# Functional crosstalk between the carrier translocase machinery and YME1 complex maintains mitochondrial proteostasis and integrity

**DOI:** 10.1101/2022.02.03.478940

**Authors:** Abhishek Kumar, Patrick D’Silva

## Abstract

The TIM22 pathway cargos are essential for sustaining mitochondrial proteostasis as an excess of these proteins leads to proteostatic stress and cell death. Yme1 is an inner membrane metalloprotease that regulates proteostasis with its chaperone-like and proteolytic activities. Although the mitochondrial translocase and protease machinery are critical for organelle health, the functional link between these complexes remains unexplored. The present study unravels a novel genetic connection between the TIM22 complex and YME1 machinery in maintaining mitochondrial proteostasis and quality control. Our genetic analyses indicate that impairment in the TIM22 complex rescues the respiratory growth defects of cells without Yme1. We further demonstrate that Yme1 is essential for the stability of the TIM22 complex and regulating the proteostasis of the TIM22 pathway substrates. Moreover, impairment in the TIM22 complex suppressed the mitochondrial structural and functional defects of Yme1 devoid cells. Notably, the functional dependence between the TIM22 and YME1 complexes remains functionally conserved from yeast to humans. Our findings suggest that excessive levels of the TIM22 pathway substrates could be one of the reasons for the respiratory growth defects of cells lacking Yme1 and compromising the TIM22 complex compensate for the imbalance in mitochondrial proteostasis caused by loss of Yme1.

## Introduction

Mitochondria are often referred to as the powerhouse of the cell due to their ability to generate ATP. However, apart from energy conversion, this organelle is also involved in diverse cellular processes, including iron-sulfur cluster production, several metabolites biosynthesis, and apoptosis (Lill and Muhlenhoff, 2005, Nunnari and Suomalainen, 2012, Kang et al., 2018). Therefore, proper mitochondrial function is paramount for cell survival and any impairment in mitochondrial health could be detrimental to cells. Mitochondria encompass a wide variety of proteins to execute these essential cellular functions, most of which are encoded by the nuclear genome, translated in the cytosol and subsequently translocated into different mitochondrial subcompartments with the help of specialized protein conducting machinery known as translocases (Baker et al., 2007, Neupert and Herrmann, 2007, Endo and Yamano, 2009, Wiedemann and Pfanner, 2017). The TOM complex is an outer membrane translocase that functions as a general entry gate for most incoming mitochondrial proteins (Dukanovic and Rapaport, 2011, Araiso et al., 2019). On the other hand, the IM contains two translocases: the TIM23 complex (presequence translocase) and the TIM22 complex (carrier translocase) (Chacinska et al., 2009, Schmidt et al., 2010, Dudek et al., 2013). The TIM23 complex mediates the sorting of proteins with presequences into the IM and matrix, whereas the TIM22 complex integrates polytopic membrane proteins into the IM (Sirrenberg et al., 1996, Rehling et al., 2004, van der Laan et al., 2010, Schendzielorz et al., 2018).

The TIM22 complex is evolutionarily conserved across the eukaryotic system, but its composition differs from yeast to humans. In yeast, the TIM22 complex is a 300-kDa insertase machinery consisting of membrane subunits Tim22, Tim54, Sdh3, Tim18, and small Tim proteins Tim9, Tim10, and Tim12 (Jarosch et al., 1996, Jarosch et al., 1997, Kerscher et al., 1997, Adam et al., 1999, Kerscher et al., 2000, Koehler et al., 2000, Kovermann et al., 2002, Gebert et al., 2011, Stojanovski et al., 2012). Tim22 forms the central core channel having a twin-pore structure, and its conserved regions play a critical role in recruiting the partner subunits to assemble the functional translocase (Rehling et al., 2003, Kumar et al., 2020). On the other hand, Tim18 and Sdh3 provide stability to the TIM22 complex, whereas Tim54 acts as an adaptor protein for small Tim chaperons docking (Kerscher et al., 2000, Koehler et al., 2000, Wagner et al., 2008, Gebert et al., 2011). The small Tim proteins assist in transferring hydrophobic client proteins from the TOM complex to the TIM22 complex (Wagner et al., 2008, Weinhaupl et al., 2021). The TIM22 pathway comprises an additional hexameric complex formed by another class of small Tim proteins, Tim8 and Tim13, required for the biogenesis of certain specific substrates, such as Tim23 (Leuenberger et al., 1999, Koehler et al., 1999). On the contrary, the human TIM22 complex is 440-kDa protein machinery containing the channel forming module Tim22 and small Tim chaperones Tim9, Tim10a, and Tim10b (functional homolog of Tim12) (Bauer et al., 1999, Muhlenbein et al., 2004, Qi et al., 2021). Further, two additional proteins Tim29 and AGK, are identified as novel membrane subunits of the human TIM22 complex (Callegari et al., 2016, Kang et al., 2016, Kang et al., 2017, Vukotic et al., 2017). However, despite having subtle changes in the structural composition, the TIM22 complex and its import pathway remain conserved from yeast to higher eukaryotes. Importantly, pathological variants in different subunits of the TIM22 complex are identified and implicated in various diseases such as mitochondrial myopathy, Sengers syndrome, and Mohr–Tranebjaerg syndrome (deafness-dystonia-optic neuropathy syndrome) (Koehler et al., 1999, Roesch et al., 2002, Kang et al., 2017, Vukotic et al., 2017, Pacheu-Grau et al., 2018). These early studies established the role of defective protein import through the TIM22 pathway in neurological and musculoskeletal disease development.

The model substrates of the TIM22 pathway include the large family of metabolite carrier proteins such as Pic (also referred to as Pic2), Aac2 (also referred to as Pet9), and Dic1 having six TM segments (Endres et al., 1999, Horten et al., 2020). The TIM22 complex also mediates the membrane insertion of specific components of the translocases (Tim17, Tim22, and Tim23) that contain four TM regions (Leuenberger et al., 1999, Horten et al., 2020, Paschen et al., 2000). Recently, mitochondrial pyruvate carrier (MPC) that possess three TM segments and Sideroflexins (SFXNs) having five TM regions were identified as unconventional client proteins of the TIM22 pathway (Gomkale et al., 2020, Rampelt et al., 2020, Acoba et al., 2021, Jackson et al., 2021). The substrates of the TIM22 pathway are highly hydrophobic and their proper assembly is critical for maintaining mitochondrial proteostasis. Thus, the turnover of these protein substrates needs to be efficient in maintaining mitochondrial homeostasis.

To maintain protein homeostasis, mitochondria harbor multiple quality control pathways such as mitochondrial dynamics, mitophagy, proteasome-assisted protein degradation, chaperones, and proteases (Ruan et al., 2020, Song et al., 2021). Amongst them, mitochondrial proteases operate at the molecular level and regulate quality control, ensuring a healthy mitochondrial pool (Koppen and Langer, 2007). Yme1 is an IM mitochondrial protease involved in the degradation of misfolded or unfolded proteins of IM, IMS, and specific OM proteins (Leonhard et al., 2000, Gerdes et al., 2012, Wu et al., 2018). It belongs to the i-AAA family of proteases that function in an ATP-dependent manner (Leonhard et al., 1999). Structurally, the YME1 complex encompasses the core component Yme1 and two additional subunits Mgr1 and Mgr3 (Dunn et al., 2008). Yme1 is a zinc metalloprotease consisting of a protease domain and an ATPase domain working hand in hand for substrate processing, whereas Mgr1 and Mgr3 act as adaptor proteins (Puchades et al., 2017). Yme1 regulates multiple arrays of the proteins, subunits of the electron transport chain such as Cox2 and Cox4, small Tim protein Tim10, and mitophagy selective receptor Atg32 (Weber et al., 1996, Stiburek et al., 2012, Wang et al., 2013, Spiller et al., 2015). Moreover, the loss of Yme1 results in abnormal mitochondrial morphology and increased frequency of mitochondrial DNA escape (Thorsness et al., 1993, Campbell and Thorsness, 1998, Cesnekova et al., 2018).

In higher eukaryotes, YME1L1, the functional homolog of Yme1, is involved in the cleavage of OPA1, a key regulator of mitochondrial fusion and fission (Griparic et al., 2007, Tilokani et al., 2018). Further, mutations in YME1L1 are associated with optic atrophy (Hartmann et al., 2016). Additionally, Tim17A, a substrate of the TIM22 pathway, was shown to be the degradation substrate of YME1L1, suggesting a possible interplay between quality control and the mitochondrial IM import machinery (Rainbolt et al., 2013). Interestingly, in yeast, Tim54, a membrane subunit of the TIM22 complex, is required to assemble Yme1 into the active proteolytic complex (Hwang et al., 2007). Furthermore, Yme1 was demonstrated to be required for quality control of Aac2 (Liu et al., 2015). Together, these findings suggest the possible existence of functional crosstalk between the TIM22 complex and YME1 machinery in regulating IM protein quality control.

The current study first time reveals a novel functional association between the TIM22 complex and YME1 machinery using *Saccharomyces cerevisiae* as a model system. Our genetic, biochemical, and cell biological analyses provide compelling mechanistic insights to highlight the functional dependence of the TIM22 complex and YME1 machinery in regulating IM protein quality control and, thus, overall mitochondrial health.

## Results

### The impairment in the TIM22 complex alleviates the respiratory incompetency of cells lacking Yme1

To understand the significance of the TIM22 complex in mitochondrial quality control, we investigated the effect of the impaired TIM22 translocase activity in cells devoid of Yme1. To begin with, we generated single deletion strains, *tim18*Δ (impaired TIM22 complex), *yme1*Δ, and a double deletion strain *tim18*Δ *yme1*Δ in W303 haploid background. Upon *in vivo* phenotypic analysis, the *tim18*Δ strain showed a similar growth pattern as wild type (WT) at all the indicated temperatures in YPD and YPG media (Figure 1A). On the contrary, the *yme1*Δ strain exhibited severe growth defects at 37°C in YPG and a mild growth sensitivity at 14°C in YPD and YPG media, in agreement with previous reports (Figure 1A) (Thorsness et al., 1993, Thorsness and Fox, 1993). The disruption of both Tim18 and Yme1 led to slight growth defects at 37°C in YPD media and 14°C in both YPD and YPG media (Figure 1A). Strikingly, *tim18*Δ *yme1*Δ strain displayed better restoration of cellular growth compared to *yme1*Δ cells at 37°C in YPG media (Figure 1A). Additionally, when we performed growth curve analysis at 37°C in YPG media, the *tim18*Δ *yme1*Δ strain showed significant growth rescue as compared to *yme1*Δ cells (Figure 1B). The identity of the deletion strains was further validated by immunoblotting analysis (Figure 1C). Moreover, the complementation of the *tim18*Δ *yme1*Δ strain with a construct expressing Tim18 under its native promoter displayed a severe growth sensitivity at 37°C in non-fermentable media, comparable to *yme1*Δ cells (Figure 1D). The Tim18 protein expression in *tim18*Δ *yme1*Δ strain was confirmed by immunoblotting (Figure 1E). Furthermore, we utilized *tim22* conditional mutants to validate the genetic link between these complexes (Kumar et al., 2020). Interestingly, the deletion of Yme1 in a *tim22* mutant background (*tim22^K127A^*) exhibited partial rescue in the growth defects of *yme1*Δ at 37°C in YPG media (Figure 1F) (Kumar et al., 2020). Taken together, these findings provide genetic evidence for a novel functional link between the TIM22 and YME1 complexes.

**Figure 1.**
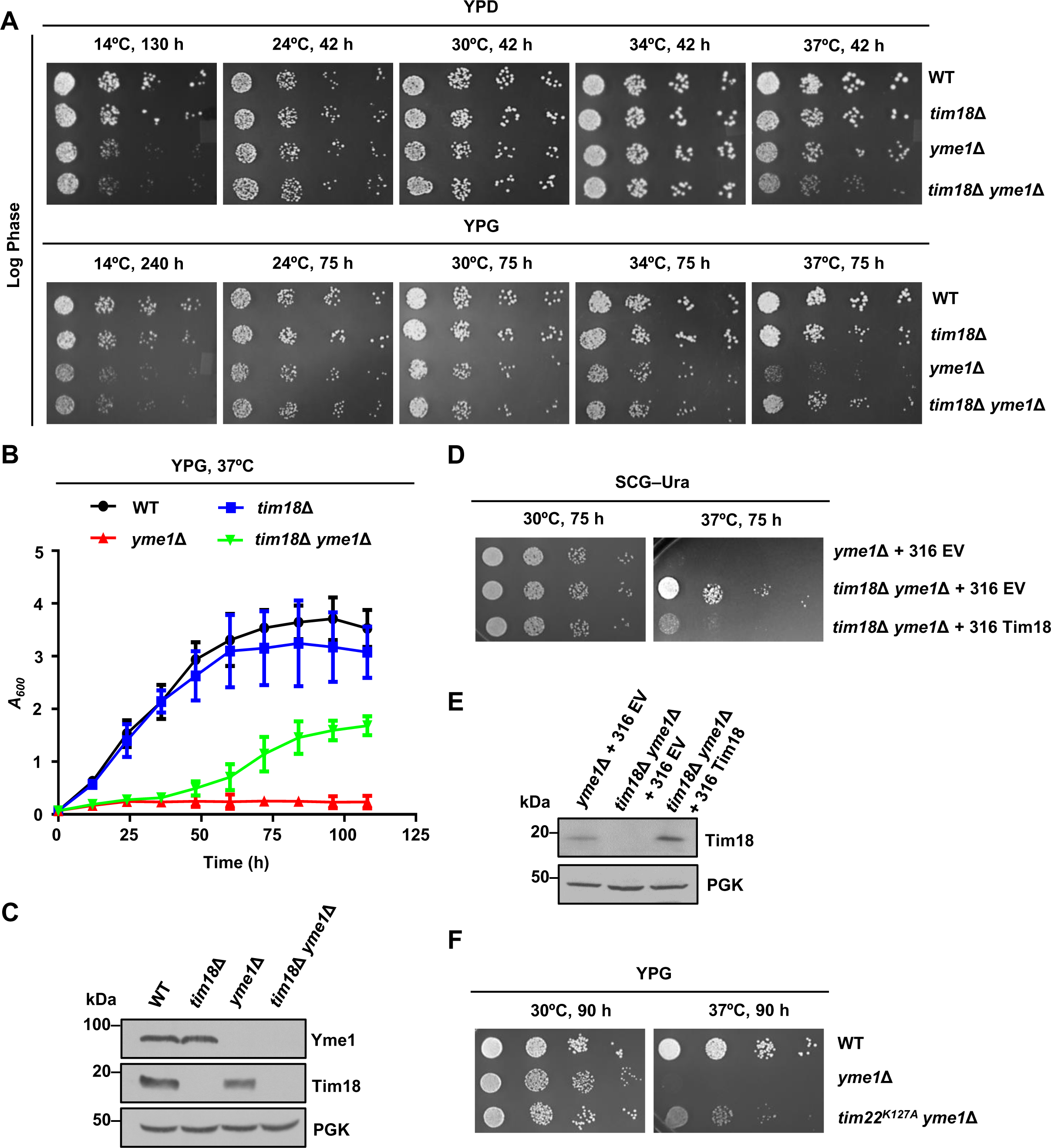
The impairment in the TIM22 complex rescues the respiratory growth defects of *yme1*Δ cells. **(A)** Growth phenotype assessment. WT, *tim18*Δ, *yme1*Δ*, and tim18*Δ *yme1*Δ yeast strains were allowed to grow up to mid-log in YPD broth at 30°C. Ten-fold serially diluted cells were spotted on the indicated media and incubated at different temperatures. **(B)** Growth curve analysis. Quantitation of the growth of WT, *tim18*Δ, *yme1*Δ*, and tim18*Δ *yme1*Δ yeast strains grown in YPG media at 37°C. **(C)** Measurement of steady-state protein levels. Whole-cell extracts of the respective strains were evaluated by immunoblotting. **(D)** Complementation of growth phenotype. *yme1*Δ and *tim18*Δ *yme1*Δ strains harboring either empty vector (EV) or Tim18 plasmid were grown to mid-log phase, serially diluted, and spotted on the SCG–Ura media. **(E)** Evaluation of Tim18 expression. Steady-state levels of Tim18 were measured in whole-cell extracts by immunoblotting. **(F)** Growth phenotype analysis. Yeast cells (WT, *yme1*Δ, and *tim22^K127A^* mutant from TM2 region of Tim22 lacking Yme1) were grown to mid-log phase, serially diluted, and spotted on the YPG media. The plates were incubated at different temperatures, and images were taken at the indicated time intervals. Data in **A**–**F** are representative of *n*=3 biological replicates.

Next, we examined if the rescue of growth defects of *yme1*Δ is specific to Tim18 loss or a consequence of impairment in the TIM22 complex. To address this, Tim18 and Tim22 were overexpressed under the *TEF* promoter and analyzed their effect on the growth of WT and *yme1*Δ strains by serial dilution assay. Remarkably, the overexpression of Tim22 severely compromised the growth of cells lacking Yme1 at all tested temperatures in both fermentable and non-fermentable media (Figure1-figure supplement 1A). At the same time, the overexpression of Tim22 did not affect the growth of the control WT strain (Figure1-figure supplement 1A). In contrast, the overexpression of Tim18 did not alter the growth phenotypes of either WT or *yme1*Δ strains (Figure1-figure supplement 1B). The Tim22 and Tim18 protein overexpression were confirmed by immunoblotting (Figure1-figure supplement 1C,D). In conclusion, these results establish that impairment in the TIM22 complex suppresses the respiratory growth defect of cells deficient in Yme1 at elevated temperature, highlighting an intricate genetic link between the carrier translocase machinery and the YME1 complex.

### Association between the TIM22 complex and YME1 machinery is critical for respiration and mitochondrial integrity

To gain insights into the respiratory growth defects of cells lacking Yme1, we first examined the oxygen consumption rate (OCR) in WT and deletion strains using Seahorse XF HS mini analyzer. Upon measurement, WT and *tim18*Δ cells did not show any significant change in the basal respiration rates (Figure 2A, Figure 2-figure supplement 1A ). However, the basal OCR was drastically reduced in *yme1*Δ strains suggesting loss of Yme1 makes the cells respiratory incompetency (Figure 2 A, Figure 2-figure supplement 1B). Remarkably, the basal OCR was rescued in the *tim18*Δ *yme1*Δ strains indicating that loss of Tim18 mitigates the respiratory incompetency of cells devoid of Yme1 (Figure 2A, Figure 2-figure supplement 1C). Additionally, we analyzed the maximal respiration capacity of WT and the deletion strains after treatment with FCCP, a mitochondrial oxidative phosphorylation uncoupler. Upon quantification, we observed that WT and *tim18*Δ cells exhibited comparable maximal respiration rates (Figure 2B, Figure 2-figure supplement 1A). Conversely, *yme1*Δ strains showed severe impairment in the maximum respiration capacity (Figure 2B, Figure 2-figure supplement 1B). Notably, the maximal respiration rate was significantly rescued in *tim18*Δ *yme1*Δ cells (Figure 2B, Figure 2-figure supplement 1C).

**Figure 2.**
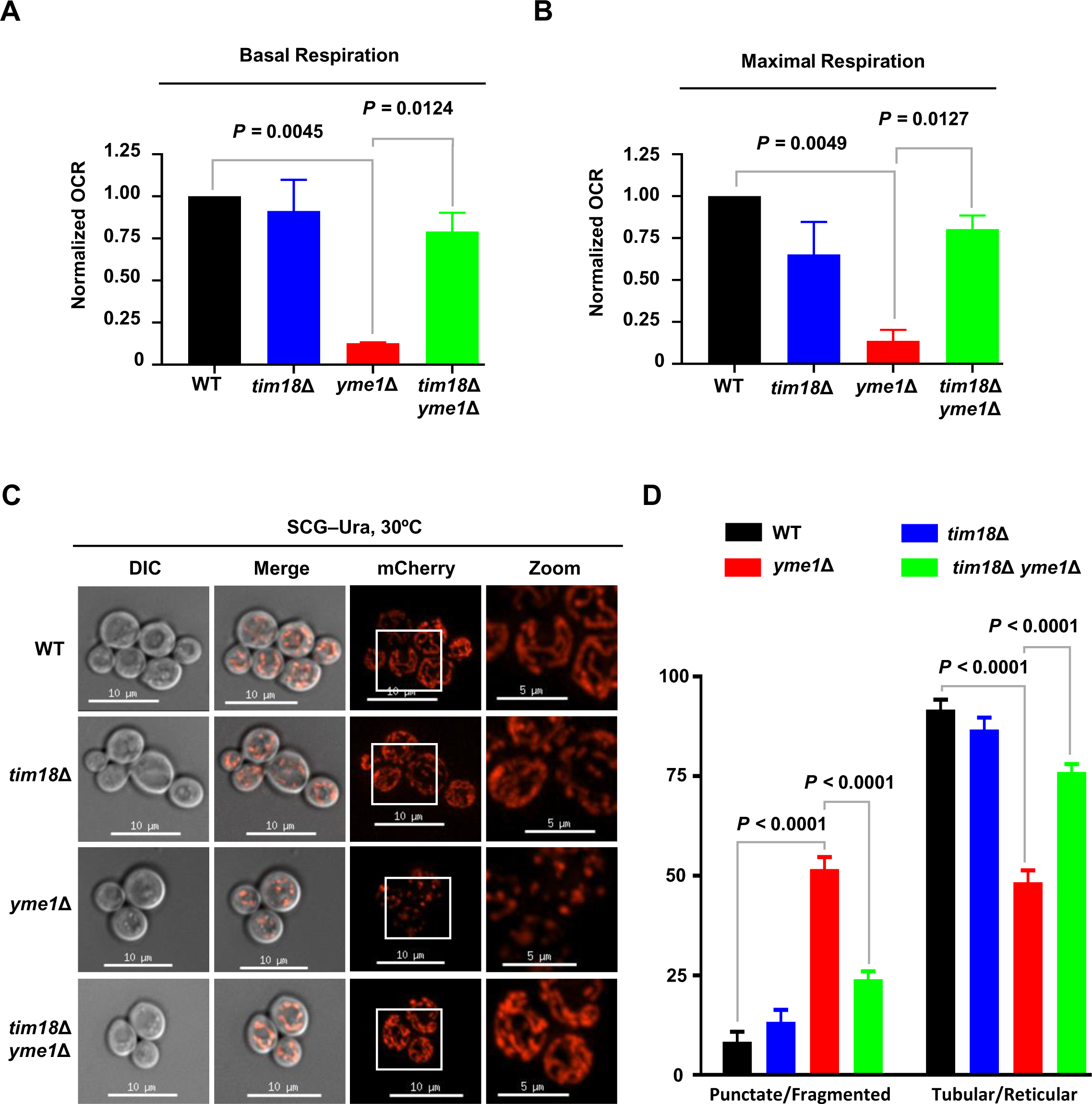
The loss of Tim18 rescues the respiration and mitochondrial morphology defects of *yme1*Δ cells. **(A,B)** Evaluation of respiration capacity. An equal number of WT and the deletion strains (*A_600_* = 0.01) were seeded in poly-D-lysine-coated Seahorse XFp cell culture miniplate to assess OCR. FCCP and rotenone/antimycin A were added one after the other to assess the mitochondrial respiratory states. Basal respiration was calculated as an average of the last OCR before the first FCCP addition subtracted by non-mitochondrial respiration capacity. Maximal respiration was measured as an average of the last OCR following the FCCP injection subtracted by non-mitochondrial respiration capacity. The non-mitochondrial respiration was calculated as an average of the last OCR after rotenone/antimycin A treatment. The fold change in basal and maximal OCR for the indicated deletion strains was calculated in comparison to WT and plotted as a bar graph. Data represent mean±s.d. of *n*=2 biological replicates with three technical replicates. Statistical significance was calculated using one-way ANOVA with Tukey’s multiple-comparison test. The representative graphs showing OCRs for WT and the deletion strains are shown in Figure 2-figure supplement 1. **(C,D)** Assessment of mitochondrial morphology. Yeast strains (WT, *tim18*Δ, *yme1*Δ, and *tim18*Δ *yme1*Δ) consisting of MTS-mCherry construct were subjected to microscopic analysis for the visualization of mitochondria. The images in all the panels were captured at identical exposures, and the mitochondrial structures from cells ≥ 50 were assessed using SoftWoRx 6.1.3 software. Boxes indicate magnified regions shown in zoom panels. Error bars in the graphs represent the s.d. in the percentage of the population. Scale bars: 10 μm and 5 μm (zoom). Data shown above are representative of *n*=3 individual experiments. Statistical significance was calculated using two-way ANOVA with Tukey’s multiple-comparison test

We next analyzed mitochondrial structures as a defect in respiration is an indication of impaired mitochondrial integrity. Since the deletion of Tim18 rescued the growth of *yme1*Δ cells (*tim18*Δ *yme1*Δ) in respiratory media, we hypothesize a possible reversal of the mitochondrial integrity through productive, functional crosstalk between the TIM22 and YME1 complexes. To validate this hypothesis, the mitochondrial morphology of WT, *tim18*Δ, *yme1*Δ, and *tim18*Δ *yme1*Δ strains were examined by microscopy. The mitochondrial network was visualized using the MTS-mCherry construct, which exclusively decorates mitochondria. WT and deletion strains expressing MTS-mCherry were grown in YPG media at the permissive temperature, and mitochondrial morphology is monitored by fluorescence imaging. Most of the WT and *tim18*Δ cells displayed a reticular or tubular mitochondrial network (Figure 2C,D). However, the *yme1*Δ strain demonstrated fragmented or punctate mitochondrial structures in agreement with the previous findings (Figure 2C,D) (Campbell and Thorsness, 1998, Campbell et al., 1994). Interestingly, *tim18*Δ *yme1*Δ cells showed enrichment of reticular or tubular mitochondrial network, suggesting that the deletion of Tim18 can rescue the mitochondrial morphological defects associated with *yme1*Δ cells under respiratory growth conditions (Figure 2C,D). Taken together, these results suggest that functional interplay between the TIM22 and YME1 complexes is crucial for mitochondrial integrity and cellular respiration.

### Crosstalk between the TIM22 complex and YME1 machinery maintains mitochondrial functions

To further delineate the importance of the genetic link between the TIM22 and YME1 complexes in mitochondrial health, we examined different mitochondrial functional parameters. Yme1 was identified in a genetic screen for isolating genes that manifest an increased frequency of mitochondrial DNA migration to the nucleus (Thorsness and Fox, 1993). We utilized mitochondrial DNA stability as one of the parameters to evaluate mitochondrial health. We used a genetic approach to examine this where the nuclear *TRP1* gene, including its control regions, was integrated into the mitochondrial genome while the endogenous *TRP1* gene was deleted (Figure 3A) (Thorsness et al., 1993). The mitochondrial *TRP1* gene, when migrated to the nucleus, becomes functional as the required transcription machinery is only present in the nucleus. The presence of *TRP1* in the mitochondrial DNA permits monitoring of the rate at which mitochondrial DNA fragments escape from the mitochondria. Upon assessment, *yme1*Δ strain demonstrated higher mitochondrial DNA transfer frequency in a time-dependent manner consistent with earlier findings (Figure 3B,C) (Thorsness et al., 1993, Cheng and Ivessa, 2010). However, the *tim18*Δ *yme1*Δ strain showed mitochondrial DNA transfer frequency comparable to WT and *tim18*Δ strains, indicating that loss of Tim18 reduces the mitochondrial DNA escape frequency of cells devoid of Yme1 (Figure 3B,C). As a loading control, an equal number of cells from WT and deletion strains were serially diluted and spotted on YPD media (Figure 3D). To further corroborate the above result, we checked for mitochondrial DNA content using mitochondrial DNA-specific dye, SYTO 18. The WT and deletion strains were grown in YPG media under permissive conditions, followed by staining with SYTO 18 and visualized by fluorescence imaging. Strikingly, the *yme1*Δ strain displayed lesser SYTO 18 fluorescence than WT and *tim18*Δ strains (Figure 3E). In contrast, the *tim18*Δ *yme1*Δ strain exhibited SYTO 18 fluorescence comparable to WT and *tim18*Δ strains, implying that the deletion of Tim18 decreases the mitochondrial DNA loss frequency of cells deficient of Yme1 (Figure 3E).

**Figure 3.**
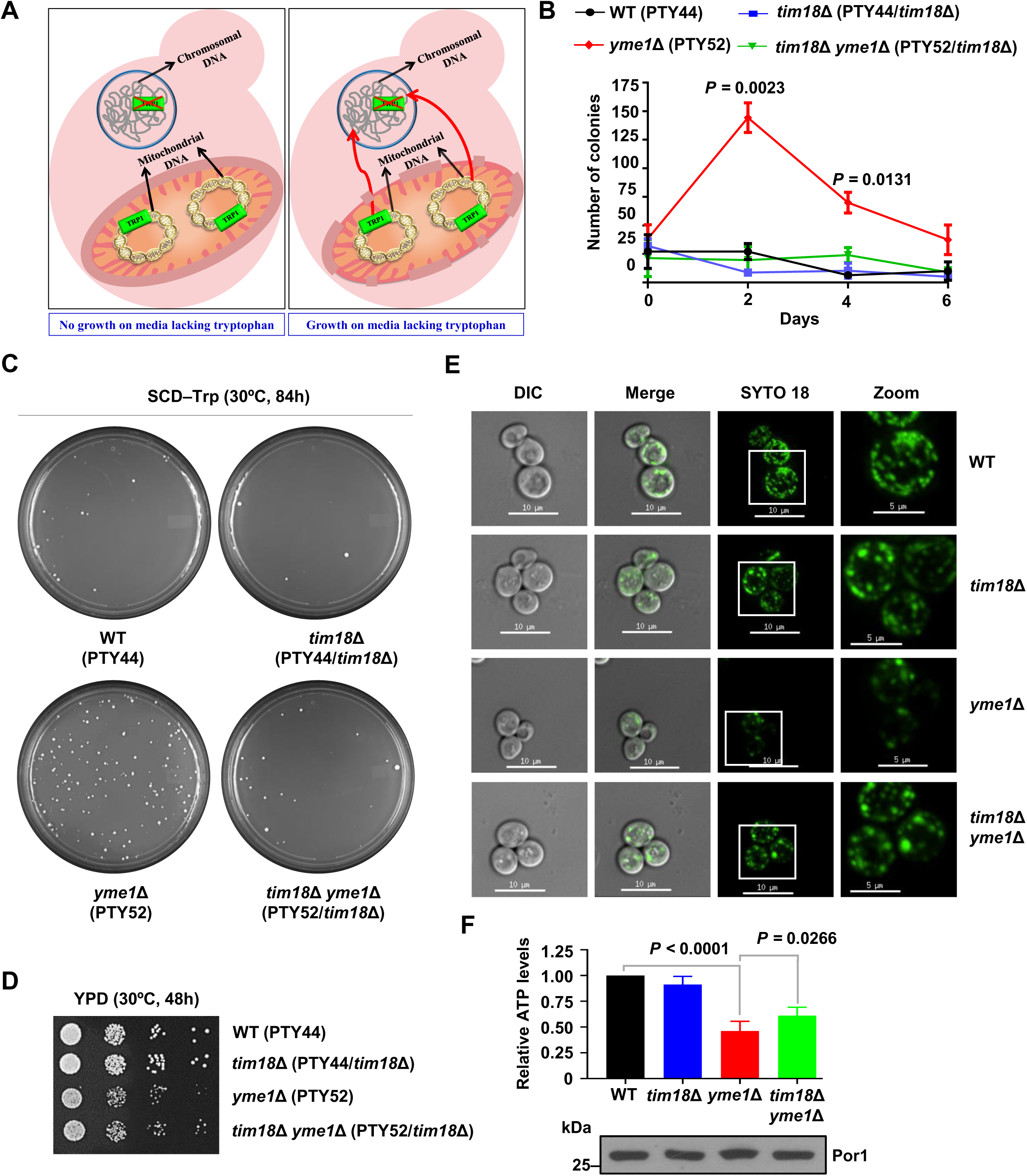
The deletion of Tim18 restores the functional mitochondrial parameters in *yme1*Δ cells. **(A)** Schematic representation of a genetic assay to determine the rate of mitochondrial DNA migration. The endogenous nuclear yeast *TRP1* gene is deleted from the genome and is inserted into mitochondrial DNA. The *TRP1* gene gets functional after it migrates from mitochondria to the nucleus since the required transcription machinery is only present in the nucleus. The migration of the *TRP1* gene from mitochondria to the nucleus allows cells to grow on media lacking the amino acid tryptophan. **(B)** Estimation of mitochondrial DNA transfer frequency. The mitochondrial DNA transmission frequency was determined by plating an equal number of cells (*A_600_* = 0.5) on media deprived of tryptophan, followed by calculating the number of colonies obtained at different time intervals and represented as mean±s.d. of *n*=3 biological replicates. Significance testing was performed using two-way ANOVA with Tukey’s multiple-comparison test. **(C)** A representative image demonstrating the number of colonies in different strains at Day 2 is shown. (**D**) WT and deletion strains were serially diluted and spotted on YPD media. **(E)** Mitochondrial DNA content analysis by fluorescence microscopy. Yeast cells grown up to the mid-log phase were stained with 10 μM SYTO 18 for 15 min, followed by mitochondrial DNA visualization using fluorescence microscopy. The images were captured by keeping all the microscopic parameters constant and analyzed using SoftWoRx 6.1.3 software. Scale bars: 10 μm and 5 μm (zoom). **(F)** Measurement of mitochondrial ATP levels. 50 μg mitochondria isolated from indicated strains were lysed and analyzed by ATP detection reagent of Mitochondrial ToxGlo Assay kit. Porin was used as a loading control. The relative ATP levels were quantitated by calculating the fold change and represented as a bar graph with mean±s.d. of *n*=3 biological replicates. One-way ANOVA with Tukey’s multiple-comparison test was used for calculating statistical significance.

We measured the ATP levels as another mitochondrial functional criterion in deletion strains to validate the above results. Upon estimation, the levels of mitochondrial ATP were drastically reduced in *yme1*Δ cells compared to WT and *tim18*Δ cells (Figure 3F). On the contrary, the deletion of Tim18 partially restores the ATP levels in cells lacking Yme1, indicating that the growth defects of the *yme1*Δ cells in respiratory media are a consequence of the impairment in mitochondrial functions (Figure 3F). Collectively, these findings signify that the functional link between the TIM22 complex and YME1 machinery plays an essential role in conserving mitochondrial functions.

### Yme1 is critical for regulating the proteostasis of the TIM22 pathway substrates

To get mechanistic insights into the significance of the functional crosstalk between the TIM22 complex and YME1 machinery, we analyzed the steady-state protein levels of the TIM22 pathway substrates in the cellular lysates of WT and deletion strains grown in YPG media at the permissive temperature. The protein levels of Dic1, Aac2, and Tim23 were observed to be similar between WT and *tim18*Δ strains (Figure 4A). However, the *yme1*Δ strain showed higher levels of Dic1, Aac2, and Tim23 than WT (Figure 4A). Remarkably, the levels of these proteins significantly reduced in the *tim18*Δ *yme1*Δ strain compared to the *yme1*Δ strain (Figure 4A). In summary, these results imply that the loss of Yme1 results in higher levels of the TIM22 pathway client proteins, and the deletion of Tim18 in cells lacking Yme1 decreases the excess levels of the TIM22 pathway substrates.

**Figure 4.**
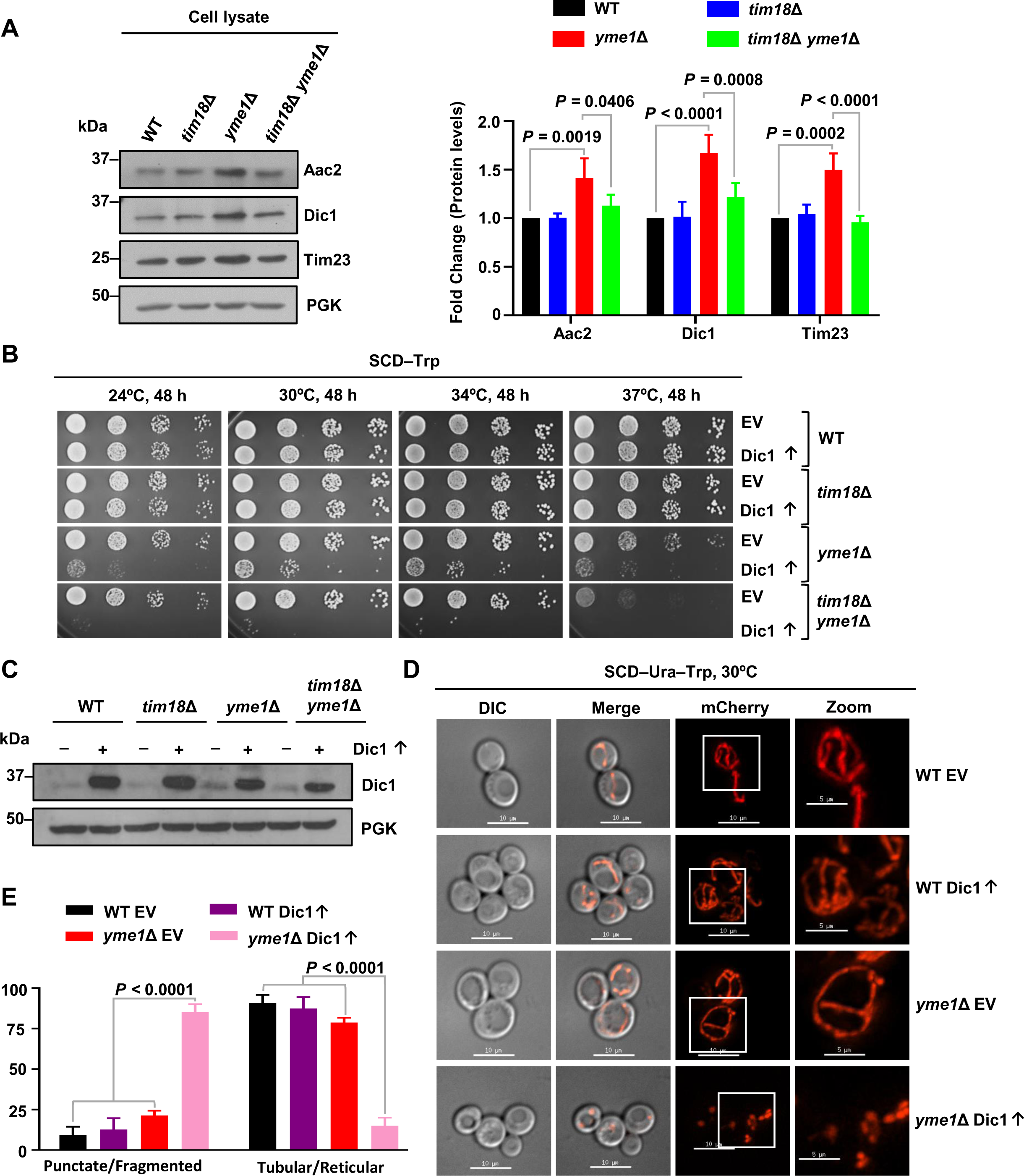
Cells lacking Yme1 exhibit excess levels of the TIM22 pathway substrates. **(A)** Analysis of steady-state protein levels of the carrier translocase machinery substrates. The protein levels of the TIM22 pathway cargos were estimated in the whole-cell lysate of WT and deletion strains grown in YPG at 30°C by immunoblotting. The protein bands were quantified using ImageJ software and plotted as fold change compared to WT protein levels. Data represent mean±s.d. Statistical significance was calculated using two-way ANOVA with Tukey’s multiple-comparison test. **(B)** Growth phenotype assessment upon overexpression of the Dic1. WT and deletion strains overexpressing Dic1 under the control of centromeric plasmid pRS414_TEF_ were serially diluted, followed by spotting on the indicated media. **(C)** Evaluation of the Dic1 overexpression. The overexpression of Dic1 was measured by western blotting in the whole-cell extracts of the indicated strains. **(D,E)** Analysis of mitochondrial morphology upon Dic1 overexpression. WT and deletion strains containing MTS-mCherry and Dic1 overexpression constructs were grown to the mid-log phase in YPD at 30°C, and mitochondrial morphology was visualized using fluorescence microscopy. The microscopic images in all the panels were taken at identical exposures, and the mitochondrial structures from cells ≥ 50 were assessed using SoftWoRx 6.1.3 software. Boxes depict magnified regions demonstrated in zoom panels. Error bars in the graphs indicate the s.d. in the percentage of the population. Scale bars: 10 μm and 5 μm (zoom). Data shown above are from *n*=3 individual experiments. Statistical significance was calculated using two-way ANOVA with Tukey’s multiple-comparison test

We next checked if the growth impairment of *yme1*Δ cells is due to the enhanced levels of the TIM22 substrates, then overexpression of these cargos should aggravate the growth sensitivity of *yme1*Δ cells. To test the hypothesis, metabolite carrier proteins (Pic, Aac2, and Dic1) and Tim23 were overexpressed followed by growth phenotype analysis. The overexpression of Dic1 did not affect the growth of WT and *tim18*Δ cells as compared to the *yme1*Δ strain that exhibited severe growth defects at all temperatures in fermentable media. (Figure 4B). Additionally, *tim18*Δ *yme1*Δ cells showed severe growth defects upon Dic1 overexpression under the same experimental conditions (Figure 4B). This signifies that the activity of carrier translocase and protease systems is crucial for handling the excess TIM22 pathway substrates. The overexpression of Dic1 was confirmed by immunoblotting (Figure 4C). Similarly, the overexpression of Tim23 considerably impairs the growth of *yme1*Δ strain at 37°C and further enhances the growth defects of *tim18*Δ *yme1*Δ strain at all tested temperatures in fermentable carbon source (Figure 4-figure supplement 1A). The overexpressed Tim23 protein levels were validated by immunoblotting in WT and deletion strains (Figure 4-figure supplement 1B). Moreover, the overexpression of more complex cargo proteins such as Pic and Aac2 resulted in the lethality of both *yme1*Δ and *tim18*Δ *yme1*Δ cells (Figure 4-figure supplement 1C,D). As a control, the overexpression of an OM protein, Tom6, or a matrix protein, Abf2, did not alter the growth of either WT or any deletion strains under similar experimental conditions (Figure 4-figure supplement 1E,F).

To correlate higher levels of TIM22 pathway substrates influence the mitochondrial integrity and thus further impair the cellular growth of *yme1*Δ cells, we examined the mitochondrial morphology in Dic1 overexpressing conditions grown in fermentable media at permissive temperature. Interestingly, WT cells overexpressing Dic1 retained their tubular mitochondrial network, whereas *yme1*Δ strain overexpressing Dic1 demonstrated further enhancement in the mitochondrial fragmentation even under fermentable growth conditions at permissive temperature (Figure 4D,E). In summary, these results suggest that excess protein levels of the TIM22 pathway substrates result in impairing the mitochondrial integrity thus leading to growth sensitivity of cells deprived of Yme1 under respiratory conditions. On the other hand, compromising the TIM22 pathway in cells lacking Yme1 reduces the higher amounts of the TIM22 pathway substrates, thereby leading to growth rescue in *tim18*Δ *yme1*Δ cells.

### Yme1 is required for the stability of the TIM22 complex

To dissect the functional link between the TIM22 complex and YME1 protease machinery, we next estimated the steady-state protein levels of different components of the carrier translocase machinery and Yme1 in isolated mitochondria from WT and deletion strains grown in non-fermentable media. Upon analysis, we found a slight reduction in Tim22 protein levels in the mitochondrial lysate of the *tim18*Δ strain at both 30°C and 37°C as compared to WT (Figure 5-figure supplement 1A, Figure 5A). This agrees with previous findings where Tim18 is demonstrated to be involved in maintaining Tim22 stability (Koehler et al., 2000, Kerscher et al., 2000). Furthermore, the levels of Tim22 and Tim18 were marginally reduced in *yme1*Δ cells at 30°C (Figure 5-figure supplement 1A). Strikingly, the levels of Tim22 and Tim18 were significantly reduced together with a minor decline in Tim54 in cells lacking Yme1 at 37°C (Figure 5A). Additionally, the *tim18*Δ *yme1*Δ strain showed a significant reduction in Tim22 levels at 30°C, whereas the Tim54 and Tim22 were considerably declined at 37°C (Figure 5-figure supplement 1A, Figure 5A). On the other hand, the levels of Yme1 protein were comparable between WT and *tim18*Δ strain at both 30°C and 37°C (Figure 5-figure supplement 1A, Figure 5A). As a positive control, the amounts of porin were examined and found to be comparable between WT and all the deletion strains (Figure 5-figure supplement 1A, Figure 5A). These results suggest that the loss of Yme1 alters the steady-state levels of different subunits of the TIM22 complex, especially at elevated temperatures.

**Figure 5.**
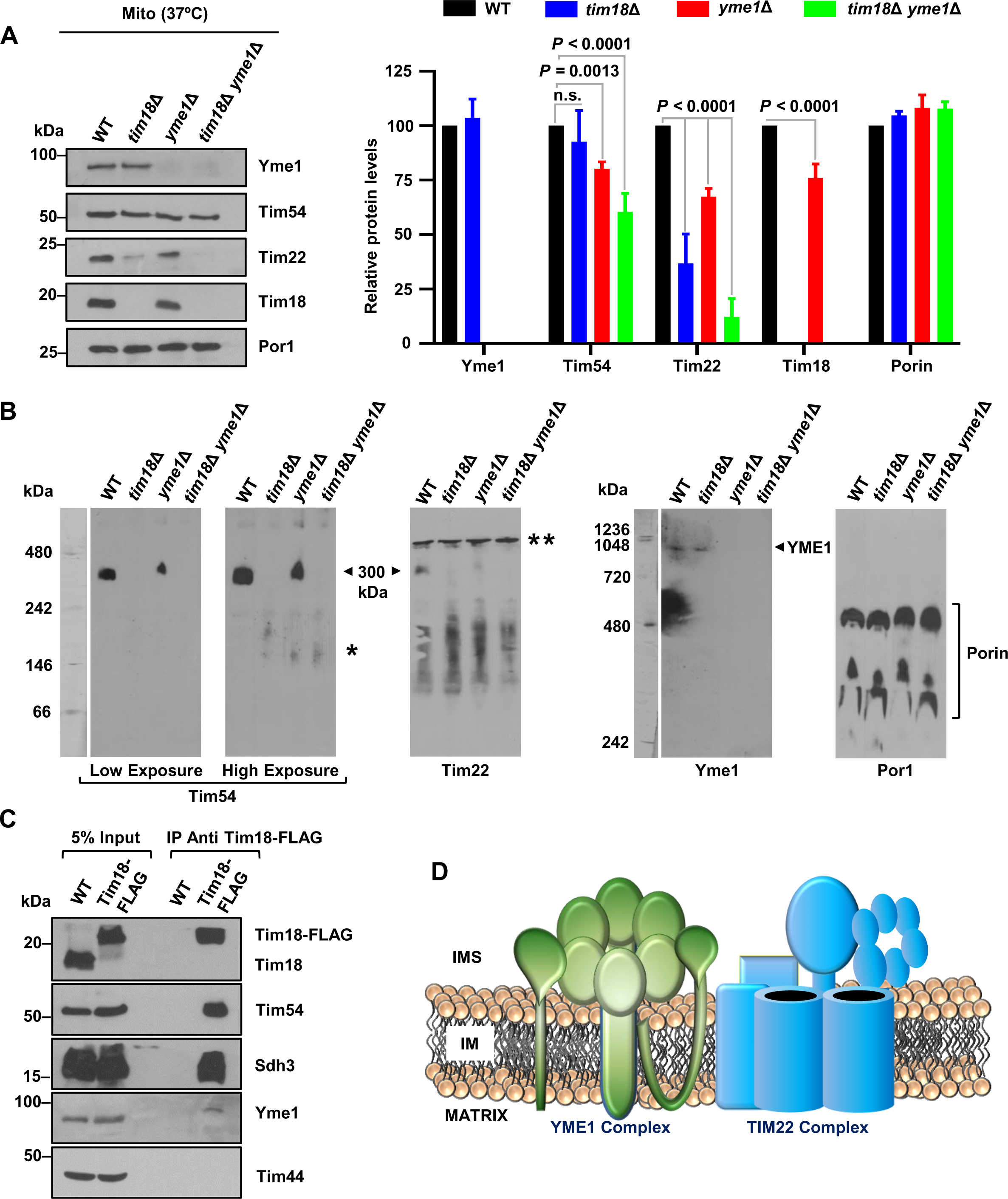
The deletion of Yme1 affects the stability of the TIM22 complex. **(A)** Estimation of steady-state protein levels of the carrier translocase machinery. 100 μg of mitochondria isolated from indicated strains grown in YPG media at 37°C were analyzed by SDS-PAGE and western blotting. Protein amounts were quantified using ImageJ software and were plotted as percentages by setting the intensities of WT mitochondria as 100%. Data indicate mean ± s.d. of *n*=3 biological replicates. Two-way ANOVA with Tukey’s multiple-comparison test was used for calculating statistical significance. **(B)** Assessment of the stability of the carrier translocase machinery using BN-PAGE. Mitochondria isolated from WT, *tim18*Δ, *yme1*Δ*, and tim18*Δ *yme1*Δ cells grown at 30°C in YPG were solubilized in digitonin buffer, and proteins were assessed by BN-PAGE followed by immunoblotting with the indicated antibodies. Arrowhead specifies the TIM22 complex (300 kDa), and the asterisk indicates possible intermediate subcomplexes. A double asterisk denotes the presence of a non-specific band. **(C)** Analysis of interaction between the carrier translocase machinery and the YME1 complex by Co-IP assay. Mitochondria isolated from WT and strains expressing Tim18–FLAG were solubilized in 1% digitonin buffer. Solubilized material was subjected to Co-IP assay using anti-FLAG-conjugated protein G Sepharose beads. Bound proteins were examined by immunoblotting using specific antibodies for the indicated proteins. 20% of the total mitochondrial lysate (input) was used as a loading control. The images shown in **B** and **C** are representative of *n*=3 experiments. **(D)** Hypothetical model suggesting the possible presence of the carrier translocase machinery and the YME1 complex in proximity to each other.

We further investigated the effect of Yme1 deletion on the stability of the TIM22 complex using blue-native PAGE (BN-PAGE). The WT mitochondria isolated from cells grown at 30°C in YPG media displayed a stable TIM22 complex at 300-kDa when immunodecorated with antibodies specific for Tim54 and Tim22 (Figure 5B). On the contrary, mitochondria isolated from *tim18*Δ *and tim18*Δ *yme1*Δ strains exhibit destabilization of the TIM22 complex (Figure 5B). This finding agrees with earlier reports where the loss of Tim18 was shown to impair the architecture of the TIM22 complex (Koehler et al., 2000, Kerscher et al., 2000). Surprisingly, the levels of 300-kDa TIM22 complex were significantly affected in cells devoid of Yme1 (Figure 5B). Furthermore, the stability of the TIM22 complex was more drastically affected at 37°C in Yme1 deficient cells (Figure 5-figure supplement 1B). This correlates well with the reduction in the levels of Tim22 and Tim18 shown above. On the contrary, the levels of the YME1 complex remained unaffected between WT and *tim18*Δ strains, suggesting that loss of Tim18 does not affect the stability of the YME1 complex *per se* (Figure 5B). Furthermore, as a control, the amounts of the porin complex were assessed and found to be comparable between WT and all the deletion strains at 30°C and 37°C (Figure 5B, Figure 5-figure supplement 1B). Together, these results highlight that deletion of Yme1 affects the stability of the TIM22 complex.

To gain more insight into the connection between these systems, we next assessed whether the TIM22 and YME1 complexes physically interact with each other. To test this, we isolated mitochondria from WT and strain expressing C-terminally FLAG-tagged Tim18 and subjected them to co-immunoprecipitation (Co-IP) using anti-FLAG prebound to protein G Sepharose beads. Upon analysis, Tim18-FLAG efficiently immunoprecipitated Tim54 and Sdh3, corroborating with previous reports (Figure 5C) (Kumar et al., 2020). Intriguingly, we observed co-purification of Yme1 along with Tim18-FLAG (Figure 5C). This is in agreement with a previous study demonstrating that Tim54 and Yme1 interact with each other (Hwang et al., 2007). Tim44, a subunit of the TIM23 complex and mitochondrial lysate from WT, were used as negative controls to determine the specificity (Figure 5C). Based on these findings, we envisioned that the TIM22 complex and YME1 machinery are in proximity to each other in the mitochondrial IM (Figure 5D).

### The impairment in the TIM22 complex rescues the respiratory growth defects of a diseased mutant in the human homolog of Yme1

A homozygous missense mutation in YME1L1, the human homolog of Yme1, has recently been reported to cause mitochondriopathy exhibiting mitochondrial fragmentation (Hartmann et al., 2016). The mutation abrogates the maturation of YME1L1, thereby leading to its rapid degradation (Hartmann et al., 2016). However, the exact pathomechanism underlying this disease progression remains poorly understood. Since we observed a functional link between the TIM22 and YME1 complex in yeast, we next investigated the significance of this crosstalk in the progression of the disease.

To address this, we first tested if YME1L1 can functionally complement the *yme1*Δ strain. Phenotypic analysis revealed that YME1L1 partially suppresses the growth defects of the *yme1*Δ strain at 37°C under respiratory growth conditions, correlating with earlier reports (Figure 6A) (Shah et al., 2000). On the other hand, the disease mutant of *YME1L1^R149W^* showed growth defects at 37°C in non-fermentable media, comparable to the *yme1*Δ strain (Figure 6A). Surprisingly, the deletion of Tim18 in the *YME1L1^R149W^* mutant showed growth rescue compared to the *YME1L1^R149W^* mutant alone at 37°C under respiratory growth (Figure 6A). Furthermore, the *tim22* TM2 mutant (*tim22^K127A^*) also partially suppresses the respiratory growth defects of the *YME1L1^R149W^* mutant at 37°C (Figure 6B). To further support the above findings we next assessed the steady-state levels of one of the TIM22 pathway cargos; Tim23 in the cellular lysates of YME1L1, *YME1L1^R149W^,* and *tim18*Δ *YME1L1^R149W^* strains grown in respiratory media at 37°C. Upon analysis, protein levels of Tim23 were elevated in the *YME1L1^R149W^* mutant compared to YME1L1 (Figure 6C,D). Conversely, the levels of Tim23 were significantly reduced in the *tim18*Δ *YME1L1^R149W^* strain compared to the *YME1L1^R149W^* mutant alone (Figure 6C,D). Together, these results establish a novel conserved genetic connection between the TIM22 complex and the YME1L1 in higher eukaryotes.

**Figure 6.**
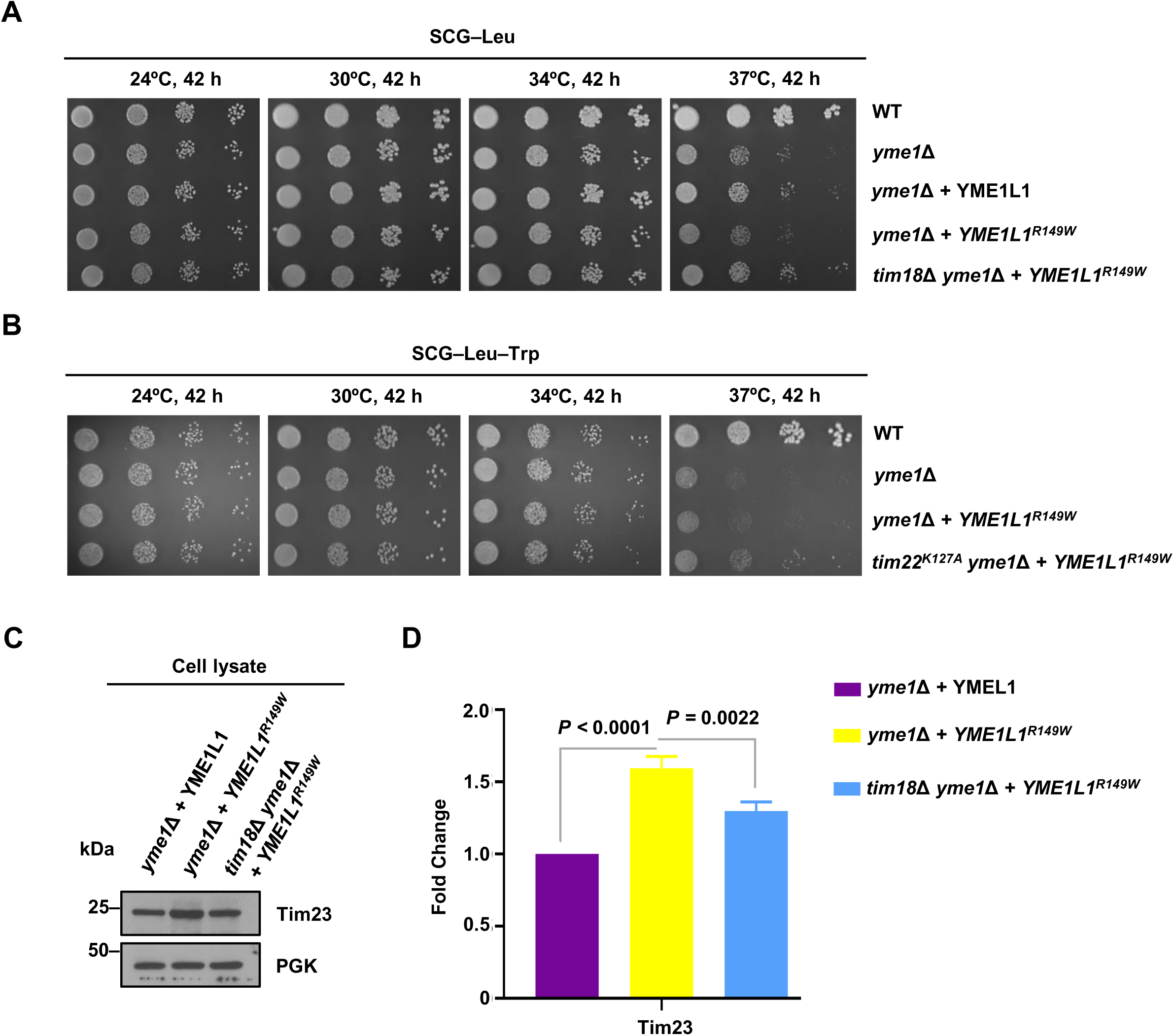
The impairment in the TIM22 complex suppresses the growth defects of a YME1L1 disease mutant under respiratory growth conditions. **(A,B)** Analysis of growth phenotype. WT and all the deletion strains were grown up to mid-log in YPD broth at 30°C, serially diluted, followed by spotting on the indicated media. The plates were incubated at different temperatures, and images were captured at an indicated time interval. Data are representative of *n*=3 biological replicates. (C,D) Assessment of steady-state protein levels of the TIM22 pathway substrates. The protein levels of Tim23 and PGK were measured in the whole-cell lysate of indicated strains grown in SCG–Leu at 37°C by immunoblotting. The protein bands were quantified using ImageJ software and plotted as fold change compared to WT protein levels. Data represent mean±s.d. of *n*=3 experiments. Statistical significance was calculated using one-way ANOVA with Tukey’s multiple-comparison test.

## Discussion

The current study highlights a functional interplay of the TIM22 complex and YME1 protease machinery in regulating mitochondrial health and integrity (Figure 7). Our genetic analyses indicate that the deletion of Tim18, a membrane component of the TIM22 complex, rescues the respiratory growth defects of cells devoid of Yme1. Additionally, one of the mutants from the TM2 region of Tim22 (*tim22^K127A^*) partially suppresses the growth defects of cells lacking Yme1 under respiratory conditions. Further, by utilizing protein overexpression, biochemical and cell biological studies, we demonstrated that the growth rescue of cells deprived of Yme1 is a consequence of impairment in the TIM22 pathway.

**Fig. 7.**
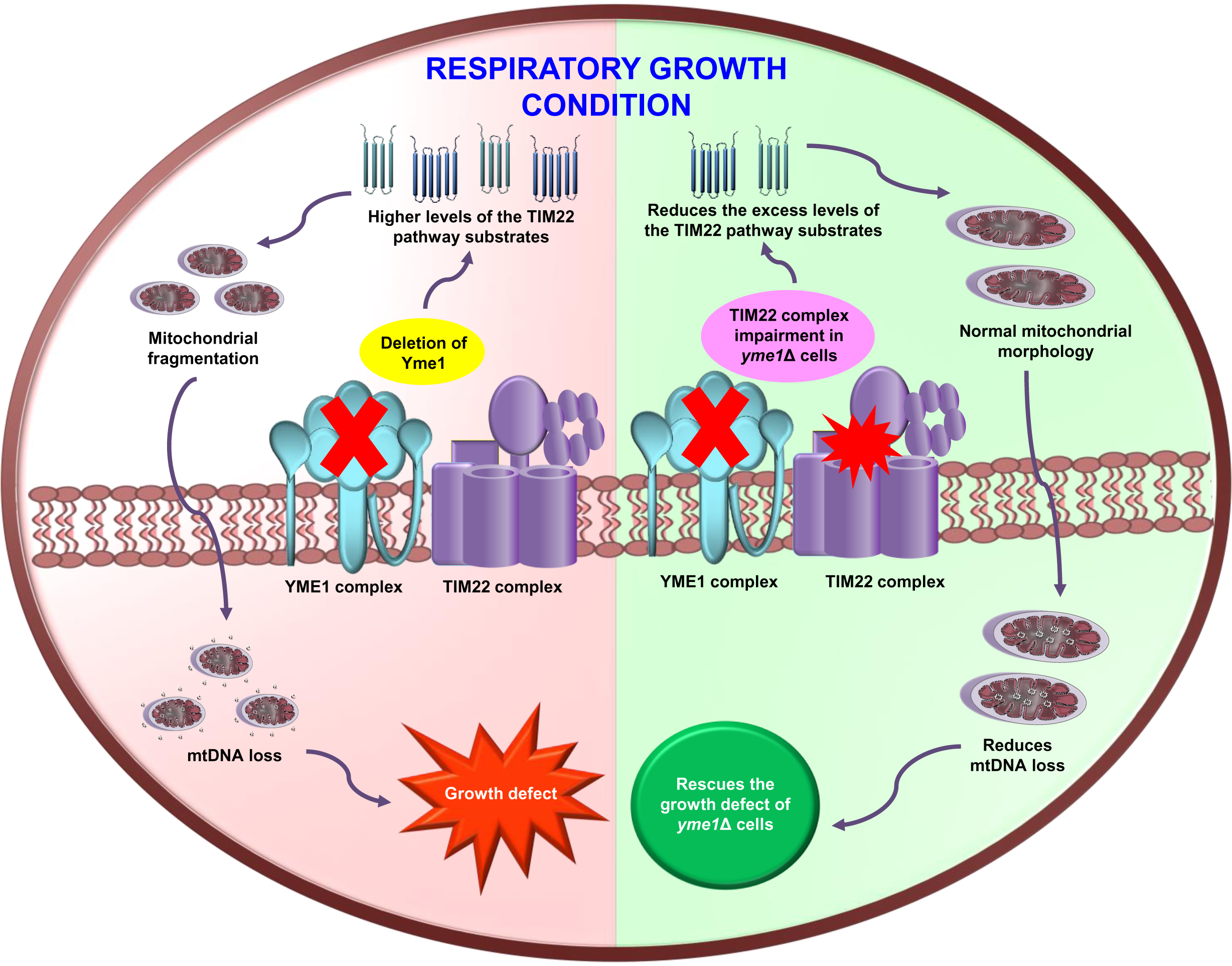
Diagrammatic illustration highlighting the significance of functional crosstalk between the carrier translocase machinery and the YME1 complex. Under the respiratory growth condition, cells lacking Yme1 exhibit growth defects and mitochondrial fragmentation due to compromised mitochondrial proteostasis and enhanced mitochondrial DNA loss. Impairment in the TIM22 complex rescues the growth and mitochondrial defects of cells devoid of Yme1 by decreasing the mitochondrial DNA loss and reducing the excess levels of the TIM22 complex substrates, thereby maintaining the mitochondrial quality control.

### Relevance of carrier translocase machinery beyond protein import

The TIM22 complex is one of the major protein translocases that performs a critical function in the biogenesis of the most intricate polytopic proteins of the mitochondrial inner membrane. However, our understanding regarding the protein homeostasis of the carrier translocase substrates is limited mainly because of a lack of studies related to the regulation of protein turnover under normal and pathological conditions. Therefore, even though the mutations in the various subunits of the TIM22 pathway are known to cause neurological disorders, the underlying mechanisms in which the levels of carrier translocase substrates are regulated under such conditions remain enigmatic.

One of the mechanisms by which cells modulate organelle health is by the action of various proteolytic machinery. Yme1 is an i-AAA mitoprotease that plays a crucial role in mitochondrial quality control by regulating the turnover of several membrane proteins. Though Yme1 is vital for mitochondrial protein turnover, cells have evolved with different adaptative mechanisms to survive in the event of its impairment or absence. Previous studies report mutations in the 26S protease subunit and gamma subunit of mitochondrial ATP synthase to repress the mitochondrial function and growth defects of *yme1*Δ cells without mitochondrial DNA respectively (Campbell et al., 1994, Weber et al., 1995). In a similar line, our study highlights that the impairment in the TIM22 complex promoted survival of Yme1 deficient cells by rescuing mitochondrial health.

The primary canonical function of the TIM22 complex is attributed to import complex IM polytopic substrates. However, recent studies highlight that the role of carrier translocase machinery is not restricted to only protein trafficking. For example, impairment in the TIM22 complex demonstrated resistance to Nde1 mediated cell cytotoxicity, implying a possible role of the protein import machinery in regulating cell death (79). Additionally, Sdh3 is a common component of both the TIM22 complex and respiratory complex, thus bridging the carrier translocase machinery with bioenergetics (24). In line with these findings, the present study provides evidence of the TIM22 complex equally involved in regulating proteostasis by working in synergy with the YME1 machinery.

### The physiological relevance of the association between import and protease machinery in regulating organellar health

The dynamic association of protein translocases with other mitochondrial modules plays a crucial role in modulating organelle homeostasis. For example, the TIM22 complex associates with porin (also known as VDAC) and mitochondrial contact site and cristae organizing system (MICOS) complex to facilitate carrier protein biogenesis (Callegari et al., 2019, Ellenrieder et al., 2019). On the other hand, the TIM23 complex coordinates with the respiratory complex to maintain the optimum IM potential required for the protein import (Wiedemann et al., 2007, Matta et al., 2020). In agreement, our finding demonstrates similar coordination between TIM22 complex and YME1 machinery being in proximity to each other to maintain mitochondrial proteostasis and organelle quality control. Notably, the interaction between carrier translocase machinery and the YME1 complex is transient since Yme1 interacted with Tim18-FLAG in a non-stoichiometric manner as observed by Co-IP. We believe that a small portion of YME1 machinery is associated with the TIM22 complex, giving rise to different pools of protease machinery to manage diverse substrates. At the organelle level, the loss of Yme1 results in punctate mitochondrial structures, enhanced mitochondrial DNA transfer, and compromised respiration capacity. Intriguingly, the defects can be suppressed by altering the TIM22 translocase activity. Hence, it is not unreasonable to believe that the IM proteostasis machinery plays a crucial role in the turnover of misassembled proteins in close connection with import machinery.

### Crosstalk between import and protease machinery regulates IM protein quality control and stress responses

Mitochondrial dysfunction is associated with several pathological states, and hence cells have evolved multiple stress-regulated pathways to maintain mitochondrial proteostasis and organelle function. Since most of the substrates of the TIM22 pathway are IM proteins with multiple TM segments, any impairment in the TIM22 complex functionality may accumulate these highly hydrophobic substrates, which were previously shown to activate mPOS (Wang and Chen, 2015). Besides, mPOS is connected to other cellular stress responses such as mitochondrial unfolded protein response (UPRmt), unfolded protein response triggered by mistargeting of proteins (UPRam), and integrated stress response (ISR) (Coyne and Chen, 2018). Such stress responses are implicated in several pathological conditions like aging, muscle degeneration, and neurodegeneration (Lin and Beal, 2006, Tatsuta and Langer, 2008, Zhu et al., 2021). In addition, the accumulation of these hydrophobic proteins in the cytosol further dampens the import of other essential cytosolic and mitochondrial proteins (Coyne and Chen, 2019). Also, overloading of the carrier proteins in humans induces cytosolic aggresomes and activates apoptotic pathways (Liu et al., 2019). Intriguingly, in our study, *yme1*Δ cells displayed higher levels of the substrates of the TIM22 pathway. Additionally, loss of Tim18 in *yme1*Δ strain reduced the accumulation of the TIM22 pathway cargos, thereby possibly suppressing mPOS and ensuring cell survival. Furthermore, since Tim18 is required to provide structural stability to the TIM22 complex, deletion of it may create a retrograde response from the organelle to the nucleus triggering anti-degenerative genes involved in UPRmt or UPRam (Wrobel et al., 2015, Fiorese et al., 2016, Labbadia et al., 2017).

Previous studies highlight that apart from stress responses, cells regulate the health of mitochondria by numerous other mechanisms, including modulating the mitochondrial protein translocation rate and its components to avoid the accumulation of misfolded proteins within the mitochondria. For example, degradation of Tim17A in higher eukaryotes limits the protein import efficiency, thereby assisting the mitochondrial proteostasis in response to stress-regulated translation attenuation (Rainbolt et al., 2013). Interestingly, our study provides evidence to show that the Yme1 is required for maintaining the stability of the TIM22 complex. We hypothesize that this might be an adaptive strategy to regulate IM protein content due to the unavailability of Yme1. Our study provides compelling evidence that in the absence of Yme1, even though the cells harbor a higher load of substrates due to partial destabilization of the TIM22 complex, the deletion of Tim18 resulted in the relieving excess of mitochondrial protein burden leading to cell survival.

### The functional link between the import and protease machinery is essentially evolutionary conserved

Conservation of critical cellular mechanisms and pathways is vital for the survival of organisms during evolution. YME1L1, the human homolog of Yme1, regulates mitochondrial dynamics by participating in OPA1 processing (Griparic et al., 2007, Tilokani et al., 2018). Mutation in YME1L1 is associated with the mitochondriopathy displaying fragmented mitochondrial structures (60). Interestingly, a mutation in the Tim22 has also been associated with mitochondrial myopathy exhibiting partially fragmented mitochondrial morphology (Pacheu-Grau et al., 2018). However, the precise pathomechanism underlying these disorders remains unclear. In the current work, our genetic analyses revealed that deletion of Tim18 or mutation in the TM2 region of Tim22 (*tim22^K127A^*) alleviates the respiratory growth defects of a YME1L1 disease mutant. Importantly, K127 residue of TIM22 has remained evolutionary conserved and is essential for the association of Tim22 with Tim18 in yeast, whereas in higher eukaryotes it is essential for Tim22 and Tim29 interaction (Kumar et al., 2020, Valpadashi et al., 2021). Further, Tim29 is required for the structural integrity and translocation of substrate proteins through the human TIM22 complex (Callegari et al., 2016, Kang et al., 2016). These results highlight that a possible functional link between the TIM22 complex and YME1 machinery is an evolutionarily conserved feature in the eukaryotic lineage.

In summary, the present study revealed a unique functional link between the TIM22 pathway and YME1 machinery in maintaining mitochondrial proteostasis and integrity. Thus, our results provide a primary platform to explore further the pathophysiology associated with YME1L1 and the TIM22 complex in humans. Therefore, the current findings are crucial for understanding how interconnection between the mitochondrial protein translocases and protease systems regulates global organelle health across the phylogeny.

## Materials and methods

### Yeast strains and genetic manipulations

The yeast strains used in this study are mentioned in the S1 Table. The details of plasmids and primers utilized in this study are described in the S2 Table. The *tim18*Δ and *yme1*Δ strains were generated by PCR-mediated homologous recombination to substitute the ORFs with a *kanMX4* and *hphNT1* selection cassette, respectively, in a W303 haploid background (91). The *tim18*Δ *yme1*Δ strain was created by deleting *YME1,* as mentioned above, in a *tim18*Δ strain. The PTY44/*tim18*Δ and PTY52/*tim18*Δ strains were made by removing *TIM18* in PTY44 and PTY52 strains following a similar approach described above. The WT *TIM18* gene from positions −700 to +859 was PCR amplified from W303 yeast genomic DNA for complementation study. The PCR-amplified products were cloned into NotI/SalI digested yeast centromeric vector pRS316. For overexpression of Pic, Aac2, Dic1, Tim23, Abf2, and Tom6, the corresponding ORFs were PCR amplified from yeast genomic DNA and cloned into the pRS414_TEF_ vector. The *TIM22* gene was PCR amplified from yeast genomic DNA and cloned into the pRS416_TEF_ vector for overexpression experiments. The *YME1L1* ORF was PCR amplified from cDNA of HEK293T cell line and cloned into pRS415_GPD_ vector for functional complementation analysis. The R149W mutation in YME1L1 was introduced by PCR-based site-directed mutagenesis using the Pfu Turbo DNA polymerase (Stratagene). The Tim22 mutant and other overexpression constructs used in this study are described in detail in an earlier report (Kumar et al., 2020).

### Media and growth conditions

Yeast cells were grown in YPD (1% yeast extract, 2% peptone, and 2% dextrose), YPG (1% yeast extract, 2% peptone, and 2% glycerol), YPL (1% yeast extract, 2% peptone, and 2% lactate, pH 5.6), SCD-Trp (0.67% yeast nitrogen base without amino acids, 0.069% Trp dropout supplement and 2% dextrose), SCD-Ura (0.67% yeast nitrogen base without amino acids, 0.072% Ura dropout supplement and 2% dextrose), SCD-Ura-Trp (0.67% yeast nitrogen base without amino acids, 0.069% Ura Trp dropout supplement and 2% dextrose), SCG-Ura (0.67% yeast nitrogen base without amino acids, 0.072% Ura dropout supplement and 3% glycerol), SCG-Leu (0.67% yeast nitrogen base without amino acids, 0.069% Leu dropout supplement and 3% glycerol), and SCG-Leu-Trp (0.67% yeast nitrogen base without amino acids, 0.069% Leu Trp dropout supplement and 3% glycerol) .

Strains harboring the *kanMX4* or *hphNT1* selection marker were selected on YPD supplemented with either 400 μg/ml G418 sulfate or 250 μg/ml hygromycin. Yeast cells were streaked and cultured on SCD containing 1 mg/ml 5-FOA to remove the *URA3*-harboring plasmid.

### Growth phenotype analysis

For spot assay, respective yeast strains were grown at 30°C in YPD or indicated media up to the mid-log phase. Cells were harvested by centrifugation (2200 ***g*** for 5 min at room temperature) followed by serial dilution and spotting on YPD, YPG, YPL, or other required minimal media. Plates were incubated at various temperatures, followed by documentation of images at appropriate time intervals.

For growth curve examination, WT and deletion strains were cultured at 30°C in YPD for 12 h. Following this, an equal amount of cells (*A_600_* = 0.07) were harvested by centrifugation (2200 ***g*** for 5 min at room temperature), washed twice with sterile water, and re-inoculated in 50 ml YPG. Absorbance was measured at 600 nm after every 12 h intervals till 4 to 5 days using a UV-visible spectrophotometer (Eppendorf).

### Whole-cell lysate preparation

Total cell protein lysates were prepared by the trichloroacetic acid precipitation method. An equal number of cells (*A_600_* = 5) were harvested by centrifugation at 16,900 ***g*** for 5 min at room temperature, resuspended in 10% TCA, vortexed thoroughly, and incubated for 30 min at 4°C with gentle vortexing at 10 mins intervals. The samples were centrifuged (2200 ***g*** for 5 min at 4°C), and trichloroacetic acid was removed completely. The resultant pellet was washed twice with acetone, followed by incubation at 42°C for 10 min. Acid washed 0.5 mm glass beads were added in 1:1 ratio (w/w) to the pellet along with 1× SDS-PAGE sample buffer (50 mM Tris-HCl, pH 6.8, 2% SDS, 0.1% bromophenol blue, 10% glycerol, and 100 mM β-mercaptoethanol) and vortexed at maximum speed for 10 times with intermittent incubation on ice. The samples were boiled at 90°C for 10 min, and the final whole cell lysate was extracted by centrifugation (16,900 ***g*** for 10 min at room temperature). Lysate volume equivalent to *A_600_* = 0.5 or 1 (depending on protein abundance) was resolved on SDS-PAGE followed by western blotting.

### Evaluation of OCR by Seahorse XF HS mini analyzer

The respiratory capacity was assessed using the Seahorse XF HS mini analyzer (Agilent Technologies). Seahorse XFp cell culture miniplate was coated with 50 μg/ml poly-D-lysine (50 ul each well) (Sigma-Aldrich) for 30 min at room temperature and then aspirated followed by air drying at room temperature for 2 h. The Seahorse XFp extracellular flux cartridge was hydrated with sterile water and incubated overnight at 37°C in a non-Co2 incubator and Seahorse XF calibrant solution (Agilent Technologies). After this, the water was removed, then added the calibrant solution to the sensor cartridge and incubated at 37°C for 1 h before measurement.

Yeast cultures of WT and the deletion strains were grown in YPD media at 30°C for 12 h, followed by subculturing in YPG media for 30°C for 8 h. The cultures were then shifted to 37°C for 8 h to induce the phenotype. For measurement, all cultures were diluted to seed an *A_600_* = 0.01 cells/well into a poly-D-lysine coated Seahorse XFp cell culture miniplate in 50 μl of Seahorse XF assay media (Agilent Technologies). For background measurement, two wells containing only assay media were included. The loaded plate was centrifuged at 300 ***g*** for 1 min at room temperature with no brakes. After centrifugation, the volume of the media was made up to 180 μl, and the loaded plate was incubated for 30 min at 37°C in a non-CO2 to facilitate the transition of the plate into the Seahorse machine’s temperature. For maximal and non-mitochondrial respiration rates, 4 μM FCCP was added to port A (Agilent Seahorse XFp cell mito stress test kit) and 1.4 μM rotenone/antimycin A in port B (Agilent Seahorse XFp cell mito stress test kit). Four readings were taken for basal respiration, whereas six measurements were recorded after FCCP and rotenone/antimycin A injections.

### Imagining of mitochondrial morphology and mitochondrial DNA by microscopy

The mitochondrial morphology was analyzed following a prior procedure (Bankapalli et al., 2015). Yeast strains were transformed with the pRS416_TEF_ MTS-mCherry construct to visualize the mitochondrial morphology. MTS-mCherry specifically decorates the mitochondria as it contains MTS from subunit 9 of ATP synthase (pSU9), thereby allowing visualization of the mitochondrial network. WT and deletion strains were grown to the mid-log phase in SCG-Ura. The cells were then harvested by centrifugation (2200 ***g*** for 5 min at room temperature) followed by final suspension in 1× PBS. The slides were prepared by placing cells (*A_600_* = 0.1) on 2% agarose pads and covering them with a coverslip. The images were acquired with a Delta Vision Elite fluorescence microscope (GE Healthcare) using a 100× objective lens by keeping a uniform exposure throughout the experiment. The excitation wavelength of 587 nm and the emission wavelength of 610 nm were used for mCherry imaging. The images were deconvolved and examined using SoftWoRx 6.1.3 software.

For mitochondrial DNA visualization, corresponding yeast cells were allowed to grow up to the mid-log phase in YPG and harvested by centrifugation (2200 ***g*** for 5 min at room temperature). The cells were resuspended in 1× PBS followed by staining with 10 µM SYTO 18 for 15 min at 30°C. After this, the cells were thoroughly washed twice with 1× PBS and placed on agarose pads. Images were captured with Delta Vision Elite Fluorescence Microscope (GE Healthcare) using a 100× objective lens. The excitation wavelength (468 nm) and the emission wavelength (533 nm) were used for monitoring the SYTO 18 fluorescence. The images were processed using SoftWoRx 6.1.3. software.

### Assessment of mitochondrial DNA transfer frequency

The rates of mitochondrial DNA transfer were determined as described previously (Thorsness et al., 1993, Thorsness and Fox, 1993). Yeast strains were grown in YPD and an equal number of cells (*A_600_* = 0.5) were harvested by centrifugation (2200 ***g*** for 5 min at room temperature). The cells were resuspended in sterile water followed by plating on a synthetic growth medium lacking the amino acid tryptophan. The number of colonies obtained was counted to calculate the frequency of mitochondrial DNA transmission.

### Measurement of mitochondrial ATP levels

ATP levels were estimated using the ATP detection reagent of the Mitochondrial ToxGlo Assay kit (Promega). 50 μg mitochondria from WT and deletion strains grown at 30°C in YPG media were lysed by two times snap freezing the mitochondria in liquid nitrogen followed by thawing on ice. The ATP detection substrate was added in 1:1 volume to the mitochondrial lysate, and the luminescence was recorded using Glomax Explorer (Promega).

### Blue native polyacrylamide gel electrophoresis (BN-PAGE)

BN-PAGE was carried out following a previously published protocol (Wittig et al., 2006). Briefly, mitochondria (2 mg/mL) were solubilized in 100 µl of digitonin buffer (1% digitonin, 50 mM NaCl, 50 mM Imidazole, 2 mM 6-aminohexanoic acid, 1 mM EDTA, pH 7.0) for 30 min at 4°C. The solubilized portion was separated by centrifugation (16,900 ***g*** for 20 min at 4°C), and 15 µl of sample buffer (2.5 µl of 5% Coomassie Brilliant Blue-G and 12.5 µl of 50% glycerol) was added per 100 µl of the supernatant fraction. Samples were run on a 6–16% gradient native imidazole PAGE gel followed by western blotting and immunodetection using Tim54 and Tim22 specific antibodies.

### Co-immunoprecipitation (Co-IP)

Mitochondria (5 mg/ml) were solubilized in 1 ml of digitonin buffer (1% digitonin, 25 mM Tris-HCl pH 7.4, 50 mM KCl, 5 mM EDTA, 10% glycerol, 1 mM PMSF). The insoluble portion was removed by centrifugation (16,900 ***g*** for 20 min at 4°C), and the solubilized fraction was then incubated with 100 µl of protein G Sepharose beads prebound to anti-FLAG antibodies. The samples were allowed to gently rotate at 4°C for 12 h followed by 3 washes with 1 ml of buffer (0.05% digitonin, 25 mM Tris-HCl, 50 mM KCl, 5 mM EDTA, 10% glycerol, 1 mM PMSF, pH 7.4). The immunoprecipitated proteins were eluted by boiling at 90°C for 10 min in 1× SDS-PAGE sample buffer. The samples were separated on SDS-PAGE and examined by immunoblotting using specific antibodies for the indicated proteins.

### Statistical analysis

The protein bands were quantified using ImageJ software. Further calculations were performed in Excel (Microsoft), and all statistical analyses were executed using GraphPad Prism 6.0 software. Significance testing was performed using one-way or two-way ANOVA and Tukey’s multiple-comparison test was used to compare different deletion strain values against WT. *P-* value of *<* 0.05 was considered significant. All indicated *n* numbers represent biological replicates.

### Antibodies and Reagents

Antisera specific for Tim22 (1:250 dilution used), Aac2 (1:250 dilution used), and Dic1 (1:250 dilution used) were gifted by Prof. Agnieszka Chacinska (Warsaw University, Poland). Antisera against Sdh3 (1:250 dilution used) and Yme1 (1:250 dilution used) were gifted by Prof. Nikolaus Pfanner (University of Freiburg, Germany). Antisera specific for Tim54 (1:1000 dilution used) antisera was gifted by Prof. Toshiya Endo (Kyoto Sangyo University, Japan. Antisera against Tim44 (1:5000 dilution used) was gifted by Prof. Elizabeth A. Craig (University of Wisconsin-Madison, USA). Antisera specific for PGK1 (1:7000 dilution used) was gifted by Prof. P. N. Rangarajan (Indian Institute of Science, Bangalore). Antibodies against Tim18 (1:2500 dilution used) and Tim23 (1: 5000 dilution used) were raised in rabbits as reported previously (Pareek et al., 2013, Kumar et al., 2020). FLAG tag antibody (1:1000 dilution used) raised in mouse (F1804) was obtained from Sigma-Aldrich and Porin antibody (1:5000 dilution used) raised in mouse (16G9E6BC4) was purchased from Invitrogen.

Yeast extract, peptone, dextrose, and agar were purchased from BD Difco™. Yeast nitrogen base without amino acids, 5-fluoroorotic acid, zymolyase, and PMSF were purchased from USBiological Life Sciences. G418, hygromycin, and digitonin were purchased from Calbiochem. NAO fluorescent dye was obtained from Molecular Probes. Other fine chemicals or reagents were purchased from Sigma-Aldrich unless not specified.

### Miscellaneous

Mitochondria were isolated following a previously published protocol (Gambill et al., 1993). Western blotting was performed using standard protocols, and the immunoblots were probed using an enhanced chemiluminescence system from Bio-Rad. The primers used in this study were manufactured by Sigma-Aldrich and sequencing reactions were carried out by AgriGenome Labs Pvt. Ltd.

## Acknowledgments

We thank Prof. Thomas Fox for providing yeast strains (PTY44 and PTY52), Prof. Toshiya Endo (Kyoto Sangyo University, Japan) for giving yeast strains (WT Tim22-316 and Tim18-FLAG/WT Tim22-316) as well as Tim54 antibodies, Prof. Elizabeth A. Craig (University of Wisconsin-Madison, USA) for Tim44 antibodies and yeast vectors (pRS316, pRS414_TEF_, pRS416_TEF_, pRS415_GPD_), Prof. Agnieszka Chacinska (Warsaw University, Poland) for Tim22, Aac2, and Dic1 antibodies, Prof. Nikolaus Pfanner (University of Freiburg, Germany) for Sdh3 and Yme1 antibodies, and Prof. P. N. Rangarajan (Indian Institute of Science, Bangalore, India) for PGK1 antibodies. We thank Dr. Sandeep M. Eswarappa for providing Glomax Explorer to analyze luminescence-based experiments. We thank Vinaya Vishwanathan and Tejashree Pradip Waingankar of the PDS laboratory for their constructive and critical comments related to manuscript and microscopy experiments. We acknowledge the flow cytometry facility of the Department of Biochemistry, Indian Institute of Science, Bangalore, India, for flow cytometry-related experiments.

This work was supported by Swarnajayanthi Fellowship from the Department of Science and Technology, Ministry of Science and Technology, India (DST/SJF/LS-01/2011–2012), Department of Biotechnology (DBT-IISC Partnership Program Phase-II (No. BT/PR27952/IN/22/212/2018) and DST-FIST Programme-Phase III [No. SR/FST/LSII-045/2016(G)] to Patrick D’Silva.

Abhishek Kumar acknowledges a research fellowship from the Council of Scientific and Industrial Research.

## Author contributions

Patrick D’Silva conceptualized the study. Patrick D’Silva acquired the funding; Abhishek Kumar and Patrick D’Silva designed the experiments; Abhishek Kumar conducted the experiments; Abhishek Kumar and Patrick D’Silva interpreted results and wrote the manuscript.

---

**Figure 1-figure supplement 1.**
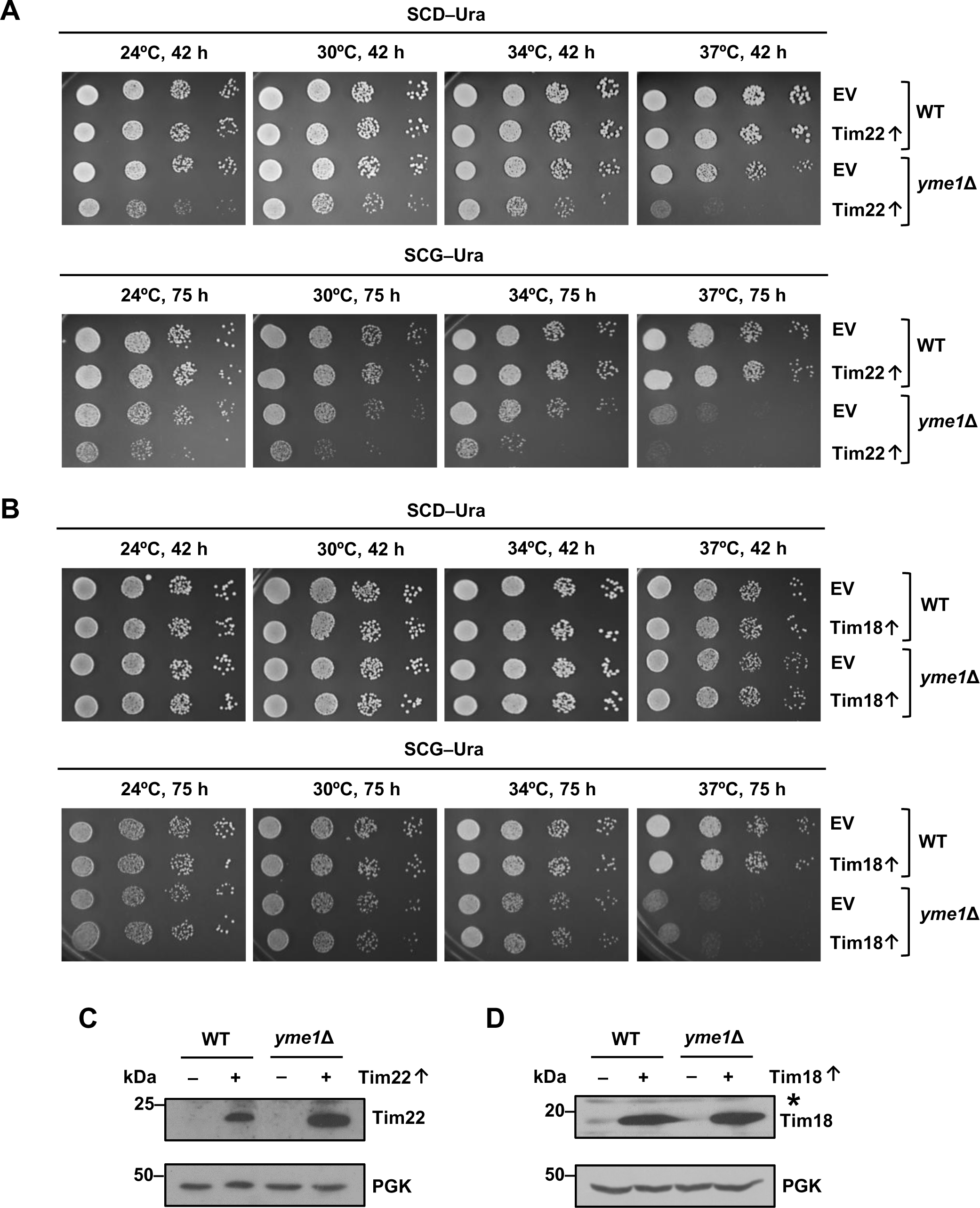
The overexpression of Tim22 aggravates the growth defects of the *yme1*Δ strain. **(A,B)** Growth phenotype analysis upon overexpression. WT and *yme1*Δ strains overexpressing either Tim22 or Tim18 under the control of centromeric plasmid pRS416_TEF_ were serially diluted and spotted on the indicated media. **(C,D)** Examination of steady-state levels of overexpressed proteins. The overexpression of Tim22 and Tim18 were examined by immunoblotting in the whole-cell extracts of the indicated strains. An asterisk indicates the presence of a non-specific band. Data in A–D are representative of *n*=3 experiments.

**Figure 2-figure supplement 1.**
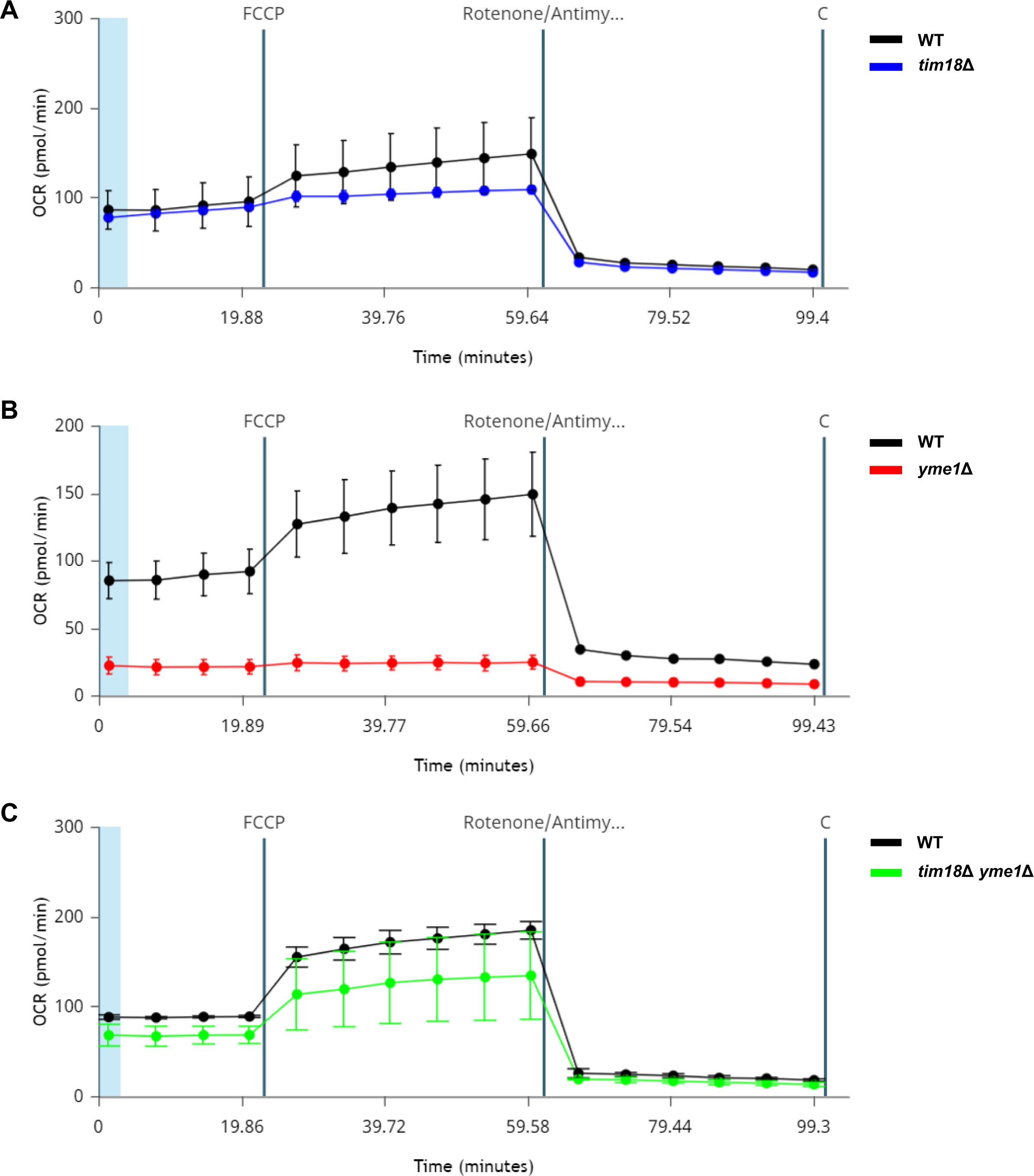
The deletion of Tim18 rescues the OCR of cells lacking Yme1. **(A-C)** Measurement of OCRs. The respiratory capacity of WT and the indicated deletion strains were determined with a Seahorse XF HS mini analyzer. FCCP and rotenone/antimycin A were sequentially added to analyze mitochondrial respiratory efficiency. Each graph represents an individual experiment showing the pattern of OCRs for WT and the deletion strains.

**Figure 4-figure supplement 1.**
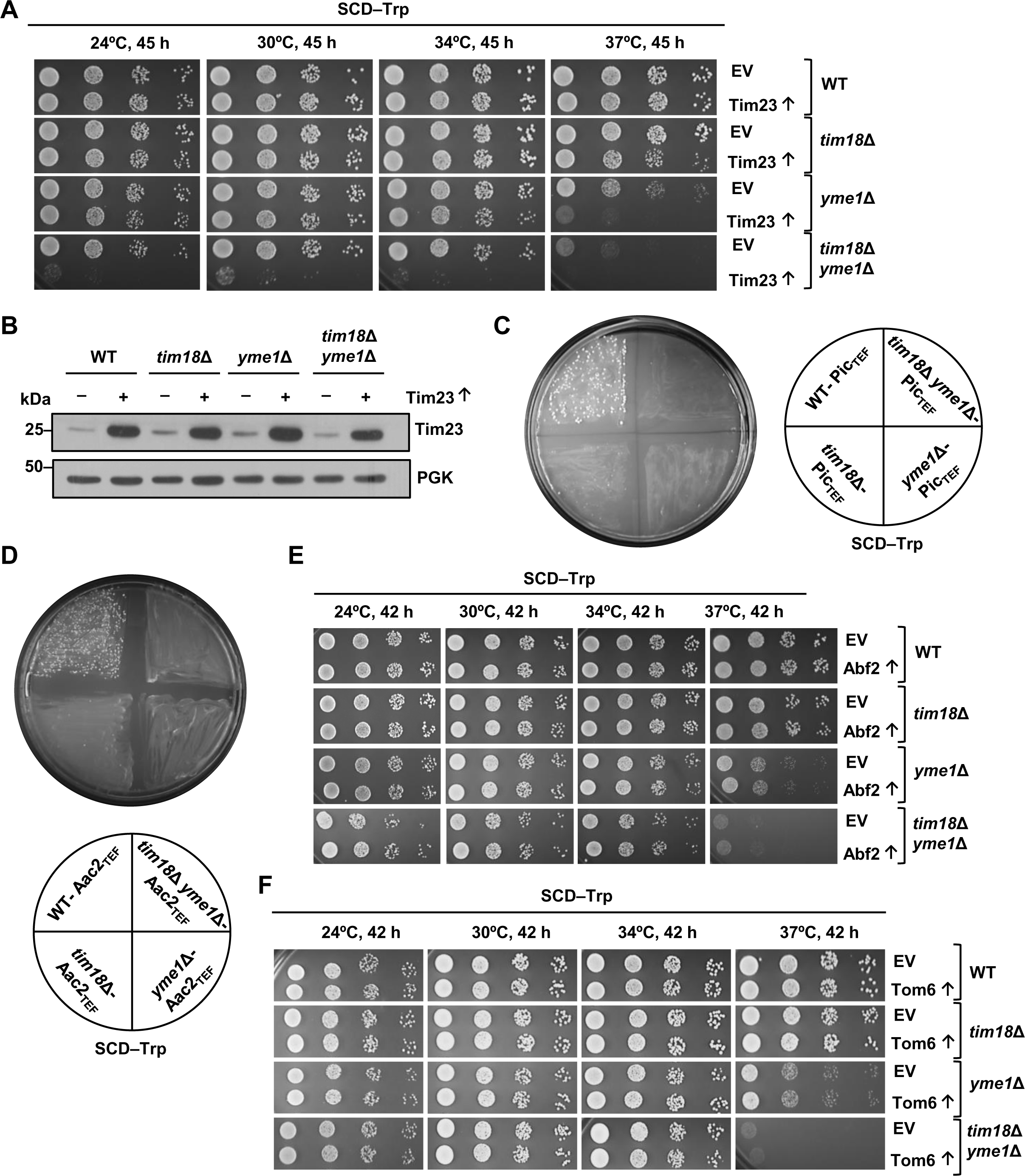
The overexpression of the TIM22 pathway substrates exacerbates the growth defects of the *yme1*Δ cells. **(A)** Growth phenotype analysis upon overexpression of TIM23. Ten-fold serially diluted WT and deletion strains overexpressing Tim23 under the control of centromeric plasmid pRS414_TEF_ were spotted on the SCD**–**Trp media and incubated at different temperatures. **(B)** Estimation of the Tim23 overexpression. The overexpression of Tim23 was examined in the whole-cell extracts of the indicated strains by immunoblotting. **(C,D)** Assessment of Pic and Aac2 overexpression on the growth of WT and deletion strains. The ORFs of Pic and Aac2 were cloned in the pRS414_TEF_ plasmid and transformed in WT and deletion strains, followed by plating on selection media (SCD**–**Trp). **(E,F)** Examination of growth phenotype upon overexpression of Abf2 and Tom6. WT and deletion strains encompassing either EV, Abf2, or Tom6 overexpressing plasmids were grown to mid-log phase, serially diluted, and spotted on the specified media. The images shown are representative of *n*=3 biological replicates.

**Figure 5-figure supplement 1.**
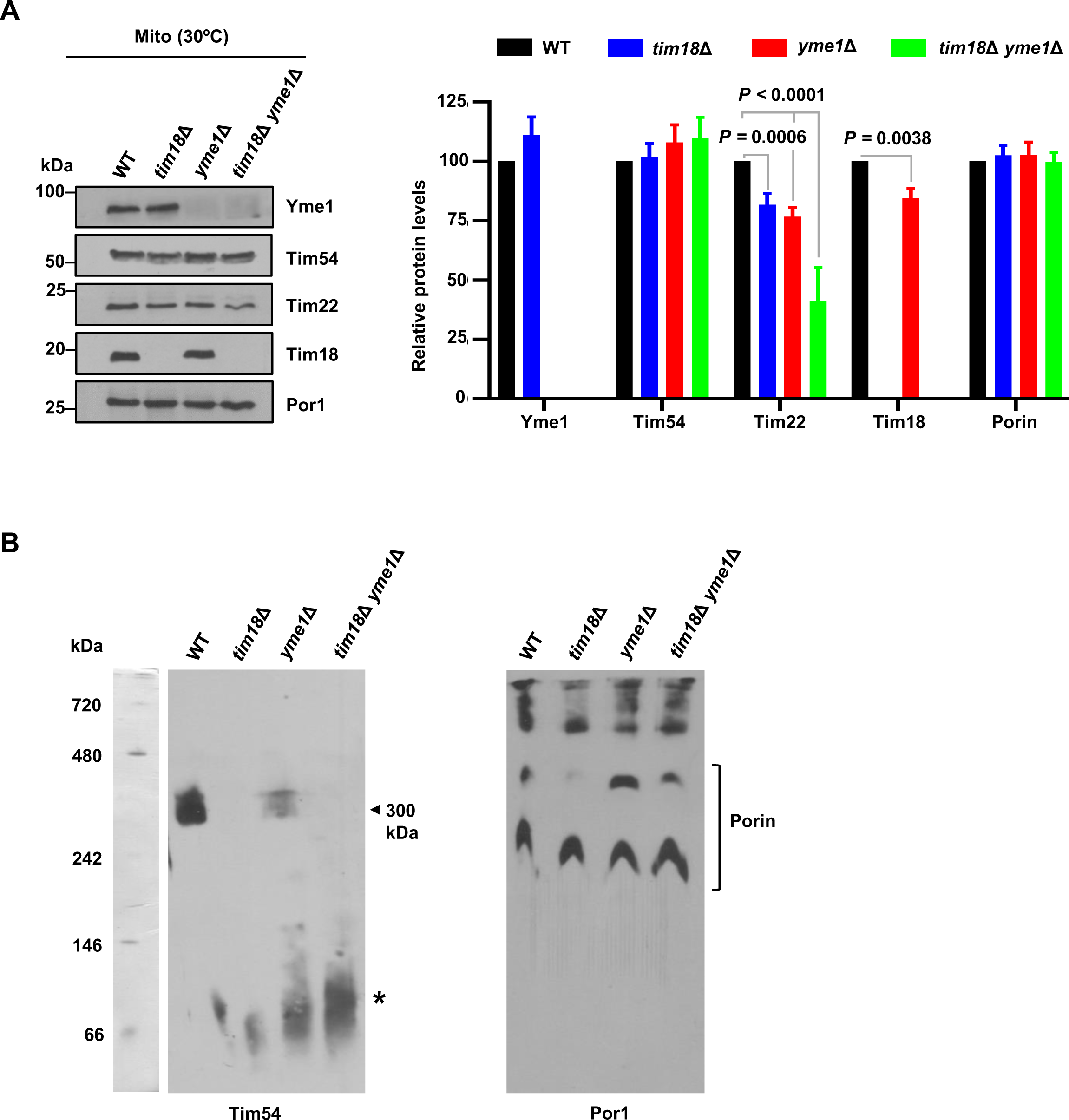
The loss of Yme1 affects the steady-state levels of the TIM22 complex components. **(A)** Measurement of the protein levels of the carrier translocase machinery. 100 μg of mitochondria isolated from indicated strains grown in YPG media at 30°C were assessed by immunoblotting. Protein amounts were quantified using ImageJ software and were plotted as percentages by setting the intensities of WT mitochondria as 100%. Data specify mean±s.d. of *n*=3 biological replicates. Two-way ANOVA with Tukey’s multiple-comparison test was used for analyzing statistical significance. **(B)** Analysis of the stability of the carrier translocase machinery using BN-PAGE. Mitochondria isolated from WT, *tim18*Δ, *yme1*Δ*, and tim18*Δ *yme1*Δ cells grown at 37°C in YPG for 24 h were solubilized in digitonin buffer, and proteins were examined by BN-PAGE followed by immunoblotting with the indicated antibodies. Arrowhead represents the TIM22 complex (300 kDa), and the asterisk indicates possible intermediate subcomplexes. Data shown are representative of *n*=3 biological replicates.

**Figure 1-source data file 1.** Assessment of steady-state protein levels.

**Figure 3-source data file 1.** Analysis of porin steady-state levels.

**Figure 4-source data file 1.** Assessment of steady-state protein levels of the carrier translocase machinery substrates.

**Figure 4-source data file 2.** Evaluation of the Dic1 overexpression.

**Figure 5-source data file 1.** Measurement of steady-state protein levels of the carrier translocase machinery.

**Figure 5-source data file 2.** Analysis of the stability of the carrier translocase machinery using BN-PAGE.

**Figure 5-source data file 3.** Evaluation of the stability of the carrier translocase machinery using BN-PAGE.

**Figure 5-source data file 4.** Evaluation of the stability of the YME1 machinery using BN-PAGE.

**Figure 5-source data file 5.** Assessment of interaction between the carrier translocase machinery and the YME1 complex by Co-IP assay.

**Figure 6-source data file 1.** Analysis of steady-state protein levels of Tim23.

**Figure 1-figure supplement 1-source data file 1.** Measurement of steady-state levels of overexpressed proteins.

**Figure 4-figure supplement 1-source data file 1.** Evaluation of the Tim23 overexpression.

**Figure 5-figure supplement 1-source data file 1.** Examination of the protein levels of the carrier translocase machinery.

**Figure 5-figure supplement 1-source data file 2.** Assessment of the stability of the carrier translocase machinery using BN-PAGE.

**S1 Table.** Yeast strains used in this study.

| Name | Genotype | Source |
| --- | --- | --- |
| WT W303 | *MATa ade2-1 his3-11, 15 ura3-1 leu2-3, 112 trp1-1 can1-100 GAL2 met2-1 lys2-2* | Hines et al., 2011 (1) |
| *tim18Δ* | *MATa ade2-1 his3-11, 15 ura3-1 leu2-3, 112 trp1-1 can1-100 GAL2 met2-1 lys2-2 tim18Δ::KanMX4* | This study |
| *yme1Δ* | *MATa ade2-1 his3-11, 15 ura3-1 leu2-3, 112 trp1-1 can1-100 GAL2 met2-1 lys2-2 yme1Δ::HphNT1* | This study |
| *tim18Δ yme1Δ* | *MATa ade2-1 his3-11, 15 ura3-1 leu2-3, 112 trp1-1 can1-100 GAL2 met2-1 lys2-2 tim18Δ::KanMX4 yme1Δ::HphNT1* | This study |
| K127A *yme1Δ* | *MATa ade2-1 his3-11, 15 ura3-1 leu2-3, 112 trp1-1 can1-100 tim22Δ::CgHIS3 [pRS314-tim22<sub>K127A</sub>] yme1Δ::HphNT1* | This study |
| WT Tim22-316 | *MATa ade2-1 his3-11, 15 ura3-1 leu2-3, 112 trp1-1 can1-100 tim22 Δ::CgHIS3, [pRS316-Tim22 ]* | Okamoto et al., 2014 (2) |
| Tim18FLAG/WT Tim22-316 | *MATa ade2-1 his3-11, 15 ura3-1 leu2-3, 112 trp1-1 can1-100 tim22 Δ::CgHIS3, TIM18-FLAG::KanMX6 [pRS316-Tim22 ]* | Okamoto et al., 2014 (2) |
| PTY44 | *MATa Lys2 ura3-52 leu2-3,112 trp1-Δ1 [rho<sup>+</sup> TRP1]* | Thorsness et al., 1993 (3) |
| PTY44/*tim18Δ* | *MATa Lys2 ura3-52 leu2-3,112 trp1-Δ1 tim18Δ::KanMX4 [rho<sup>+</sup> TRP1]* | This study |
| PTY52 | *MATa Lys2 ura3-52 leu2-3,112 trp1-Δ1 yme1-Δ1::URA3 [rho<sup>+</sup> TRP1]* | Thorsness et al., 1993 |
| PTY52/*tim18Δ* | *MATa Lys2 ura3-52 leu2-3,112 trp1-Δ1 yme1-Δ1::URA3 tim18Δ::KanMX4 [rho<sup>+</sup> TRP1]* | This study |

**S2 Table.**
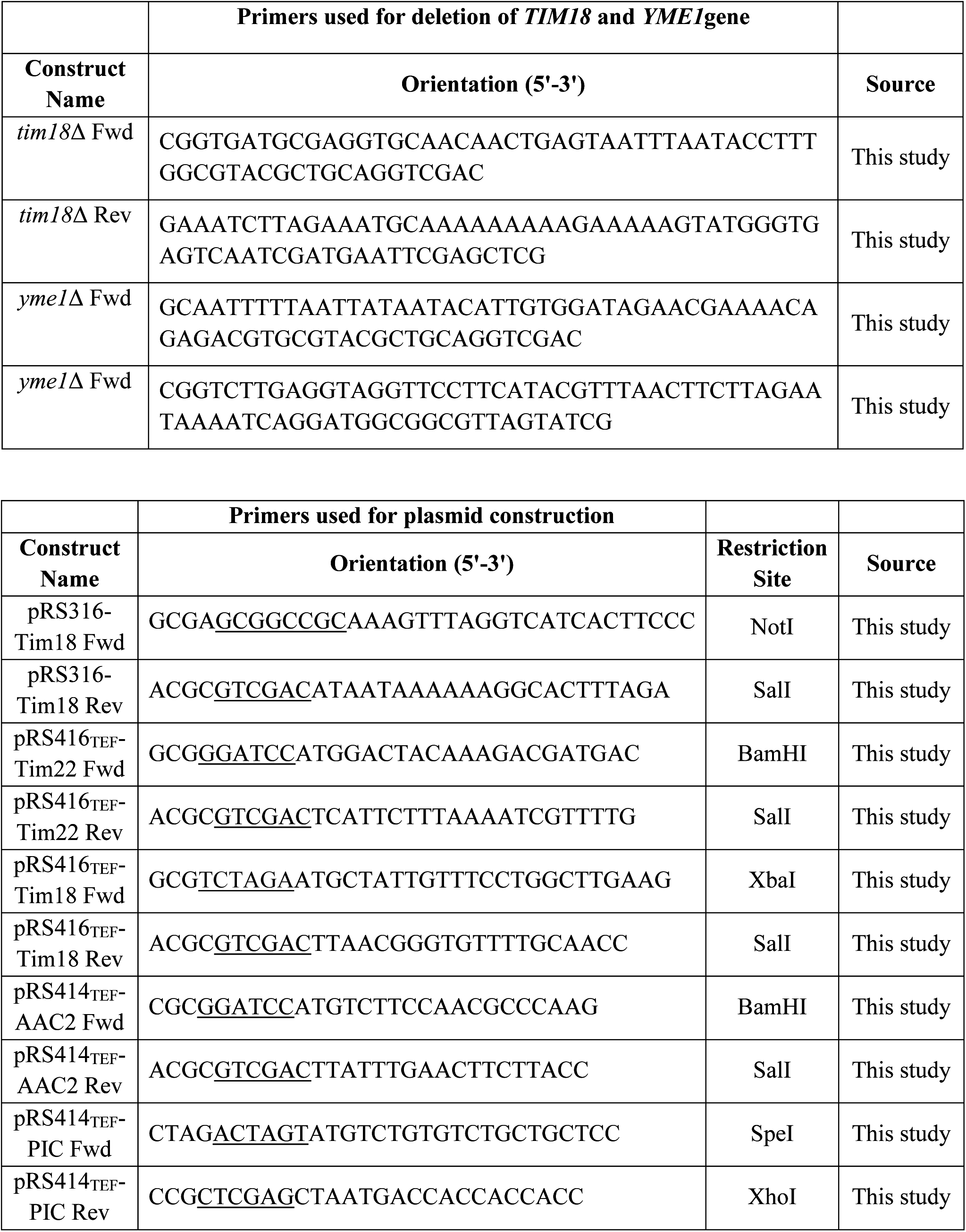
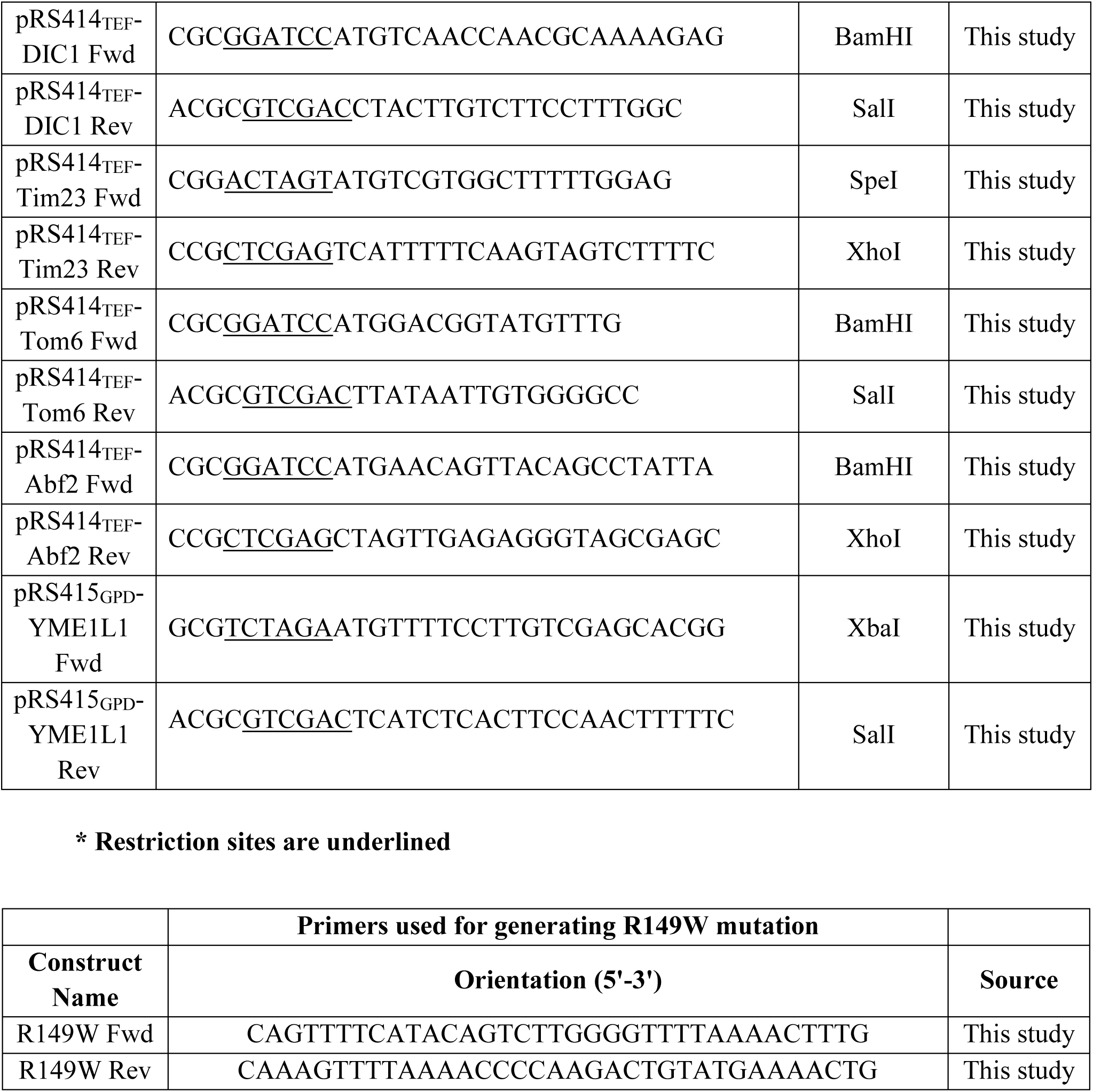
Plasmid constructs and primers used in this study.

